# Prediction of protein functions using Semantic Based Regularization

**DOI:** 10.1101/2024.06.20.599881

**Authors:** Giovanna Maria Dimitri

## Abstract

In this work, done in collaboration with Prof. Michelangelo Diligenti (department of Engineering and Mathematics, University of Siena) we present the use of Semantic Based Regularization Kernel based machine learning method to predict protein function. We initially build the protein functions ontology, given an initial list of proteins. We subsequently performed predictions, both at individual and at joint levels of functions, introducing and adding to the learning procedure ad-hoc first order logic rules. Experiments showed promising performances in using logic rules within the learning process for the sake of bioinformatics applications.

## 1 Introduction

Protein-Protein Interaction (PPI) and Protein Function (PF) predictions have a central role in biological research, since they are largely used in many fields such as drug and enzyme design. Associating a function to each protein would allow a significant advance in the understanding of biological processes and protein interactions, shedding light on disease development and aging. Unfortunately, a manual generation of a database where to each protein are associated the corresponding functions, is not feasible. For this reason, machine learning techniques are widely used to predict PFs, due to the scalability properties typical of these methods. However, prediction accuracy is often an issue when using machine learning techniques in bioinformatics. First of all, the intrinsic original nature of the problem is sometimes neglected, when trying to fit biological data into a machine learning framework. Secondly, machine learning approaches have been usually applied in one single database, thus neglecting the correlation between information stored in distinct biological databases. However combining information coming from different databases is usually a quite challenging task. In fact, biological databases (such as UniProt (3), BioGRID (5) or KEGG (6)) are often semi-manually compiled by biological experts, frequently according to incompatible schemes or discordant definitions of the same biological concept (such as gene function), making automatic interoperability and integration difficult. In an effort to lessen this problem, recently researchers have developed and applied a number of ontologies to formally define the database semantics, culminating with the OBO initiative (7). These developments try to lower the difficulties in performing automatic reasoning (i.e. automatic compilation of entries and consistency enforcement), over distinct, yet correlated, resources. Semantic Based Regularization (SBR) is a state-of-the-art statistical relational learning method, performing collective classification over attribute value representations using weighted first-order-logic (FOL) rules (2). This is the reason why its application to bioinformatics tasks gives very good results. In fact it is possible to add biological knowledge to the learning process through the introduction of FOL rules. This successful characteristic of SBR has been already analysed in the prediction of protein-protein interaction in (8) and confirmed in this work, showing the capability in predicting protein-functions. In our work we started from building the protein functions ontology of our original list of proteins. Then, we performed predictions either of individual levels of functions and of joint proteins functions levels, with the introduction of ad-hoc FOL rules. In performing experiments interesting results were obtained, in particular shedding light on the presence of a correlation between interacting proteins having the same functionalities.

## 2 Kernel Machines

Kernel methods, in machine learning, are a class of algorithms widely used for pattern analysis, whose best known example is the support vector machine (SVM) (9). Pattern analysis, in fact, aims at discovering possible regularities that data may hide. There are many different types of patterns that can be discovered, such as *exact patterns* (motions of planets), *complex patterns* genes in (DNA sequences) or *probabilistic patterns* (market research). Patterns detections gives the possibility to understand the regularities behind data organization and so making predictions for the future, and this is the reason why studying automatic detection of patterns in data is extremely important (10).

Pattern analysis algorithms must have the following features:

- Computationally efficient: the running time must be polynomial, in the size of the data, often of a low degree.
- Robust: able to handle noisy data.
- Statistical stability: able to distinguish between chance patterns and those characteristic of the data analysed.

Nowadays machine learning tasks involving pattern analysis such as classification, clustering or regression are widely approached using kernel methods. These, in fact, are characterized by a good generalization performance on many datasets. Moreover just a few number of parameters have to be adjusted during the learning process, simplifying their usage (11).

Machine learning methods can be of two fundamental types: linear or non-linear. The first ones are well known for their optimality and computational efficiency, whereas the second ones are characterized by their potential classification power. Kernel based methods tie together these two characteristics, through the mapping of the input patterns into a high dimensional features space, leaving parameter optimization linear (8). It can be shown using the representer Theorem (12), that many classes of problems admit solutions using kernel expansions, that is:

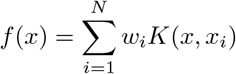

Where:

- *x* is the representation of the pattern
- *K* (*x, x*_*i*_) = ⟨Φ(*x*), Φ(*x*_*i*_)⟩ is a kernel function
- Φ(.) is a mapping from the input space to the feature space

The main idea standing behind this definition, is that the kernel function measures the similarity between pairs of instances, and *f* (*x*), that is the prediction of a novel instance, is computed as a weighted similarity to instances of the training set *x*_*i*_.

At this point it is easy to understand that the main problem is the optimization of the weights *w*_*i*_. Such an optimization problem can be stated in various ways. Suppose that pattern *x*_*i*_ has as desired outputs *y*_*j*_ ∈{ −1, +1}. Moreover *w* = [*w*_1_, …, *w*] is a vector containg the machine parameters and *G* is the gram matrix, where the (*i, j*) element is defined as: *G*_*ij*_ = *K*(*x*_*i*_, *x*_*j*_).

At this point, if we define ‖*f* ^2^‖ = *w*^*t*^ *Gw*, then we can write as cost function:

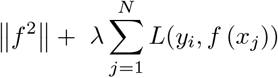

This function can be reduced to the formulation of hard margin *l*_2_SVMs if *L*(.) is the hinge loss and *λ*→ ∞ (8). Cost functions can be different depending on the problem to be solved.

### 2.1 Kernel Methods in Bioinformatics

Kernel methods have been largely used for bioinformatics tasks such in (13). In the following sections we will briefly describe some tasks in which bioinformatics have been applied.

#### 2.1.1 Identification of remote homologous proteins

At the beginning of 1990s, traditional methods were applied to discover such homologous sequences such as:

- Methods based on the similarity between couples of proteins: for example the method by Smith and Waterman (14) or heuristic methods such as BLAST (15) or FASTA (16)
- Methods based on statistical and probabilistic models: for example Hidden Markov Models (17)
- Methods based on multiple alignments comparing sequences present in databases: PSI-BLAST (18)

At the beginning of 2000s, machine learning techniques, and in particular kernel methods, started to be applied to bioinformatics tasks. In homologous proteins predictions, there were a number of methodologies applied in order to attack bioinformatics problems, such as:

- SVM Fisher kernel: this is a new method (19), called also the Fisher kernel method, used for detecting remote protein homologies. The experiments performed with this method showed good performances in classifying proteins domains by SCOP superfamily. In practice this method is a variant of support vector machines using a new kernel function. This kernel is derived from a Hidden Markov Model and the general idea standing behind this method is to use also negative examples (that is sequences that are not known as part of a specific protein family). Starting from this assumption, the HMM is trained and a gradient vector is built based on the HMM estimated parameter. At this point, the SVM is trained using such parameters. Further details could be found in (19).
- Composition kernel: proteins are considered as frequency vectors of letters. This letters can belong to six different alphabets: amino acids, secondary structure, hydrophobicity, Van der Waals volume and polarity. Support Vector Machines (SVM) and Neural Network learning methods are used as base classifiers. Further details are available in (20).
- Motif kernels: motif kernels are an extension of composition kernels. Features are composed by a motif of pre-existing databases. There are two main contributions in which motif kernels are applied. In (21), for example, the dictionary is a library of motifs and the similarity function is a matching function. Motifs were obtained by the database BLOCKS and organized in a 10.000 dimensional vector. This gave better results than SVM-Fisher kernel. In the second experiment described in (22) the database used was eBLOCKS, and 50.000 motifs were used. That is why special data structure to calculate the kernels were used.
- Kernel based on the couple comparison: the main idea standing behind these kernels is the assumption on the molecular evolution based on mutations and small insertions and deletions. The algorithm described in (23) is composed by two fundamental steps:
  1. Computation of the alignments score among all the pairs of proteins (matrix A of alignments)
  2. Each row of A represents a certain protein in the feature space: a standard kernel function can therefore be applied to compute the similarity among protein couples (also called empirical kernel map) computation of the kernel matrix K. The problem can be computationally complex, but it is possible to reduce complexity using BLAST instead of Smith Waterman algorithm for step 1.
- Spectral Kernels or Kernels for strings: these kernels operate on strings defined as symbols sequences, not necessarily of the same length. Spectral kernels measure the similarity of pairs of strings: having two strings a and b the more similar they are, the higher the value of a string kernel *K*(*a, b*). These types of kernels are used with domains composed by sequences of data which have to be clustered or classified, for example in text mining and gene analysis. In a sense, they can be seen as a generalization of composition kernels, and they have been widely applied for protein sequence prediction (24).

#### 2.1.2 Functional classification of genes and proteins

The functional classification of genes and proteins has been analysed with various methods for example based on the promoter region analysis, on the function proteins prediction based on phylogenetic profiles, on the subcellular localization prediction of proteins or on the classification through binary features obtained by multiple alignments (13).

Regarding the promoter region analysis, for example, in (25) the upstream untranslated gene region are used for extracting information to make predictions about their transcriptional regulation. The method for classifying genes is based on motif-based Hidden Markov Models (HMMs) of the correspondent promoter regions. HMMs are constructed using sequence motifs discovered in yeast promoters and HMMs also include parameters describing the number and relative locations of motifs within each sequence. Each model so constructed gives as result a Fisher kernel for a support vector machine that is usable to predict the classification of unannotated promoters. This method was adopted on two classes of genes from the budding yeast *Saccharomices Cervisiae*. The results obtained showed that the additional sequence features captured by the HMM helped in correctly classifying promoters.

An interesting example concerning the functional classification of genes and proteins is discussed in (26). Indeed in this case the protein function is predicted using the phylogenetic information contained in the protein sequence. The protein phylogenetic profile, in fact, is represented through a bit string. Each bit indicates whether or not there is an homologous gene in a given species, therefore it represents a part of the species evolutionary history. The profiles will be then mapped into a high dimensional feature vector space that incorporates phylogenetic information and in (26) is provided an algorithm that permits to compute efficiently the inner product in that space, called tree kernel. The tree kernel can be used by any kernel based analysis method for classification or data-mining of phylogenetic profiles. In the paper it was shown how using the tree kernel with an SVM, gives much better results than the SVM with a naïve kernel that contains no information about the phylogenetic history of the protein sequences analysed. Moreover the authors performed a Kernel Principal Component Analysis of the phylogenetic profiles showing the sensitivity of the tree kernel to evolutionary variations.

#### 2.1.3 Pattern recognition in biological sequences

In these cases kernel machines have been extensively applied for many tasks such as prediction for the translation starting site, prediction of splicing sites, prediction of subcellular localization or prediction of secondary protein structure.

For example in (27) the secondary protein structure is predicted, using an SVM approach with a Gaussian Kernel and a sliding window on the protein sequences. Another interesting contribution is (28) where SVMs are applied to perform translation initiation sites (TIS) prediction. The task of finding TIS is modelled as a classification problem. To incorporate biological prior knowledge they make use of an appropriate kernel function, using a fixed length window to identify genes.

In (29) starting sites of introns are predicted using a naïve Bayes model in conjunction with an SVM approach. This helps in understanding new features to be used for such prediction. They use a fixed length window, with a 4 bit codification that helps in finding the most significant position.

#### 2.1.4 Methods for integrating data biologically heterogeneous

A certain biological entity can be characterized in many different ways. An important issue in bioinformatics is to understand how these different structured data can be integrated for the sake of prediction. For example a gene can be associated to its sequence, the codifying protein sequence, the protein structure, the similarity with other proteins, the mRNA levels associated to the genes or the map of interactions with other proteins. How can all these heterogeneous data be integrated and associated to a certain gene?

Through kernels different data can be represented in a homogenous way (kernel matrix). There are various types of integration based on kernels:

- Initial integration: simple vector concatenation.
- Intermediate integration: kernels computation done separately and then they are added together. An interesting example of this is (30).
- Late integration: kernels and discriminative functions are computed separately and then integrated in a second step.
- Intermediate weighted integration: the added matrix is weighted.

#### 2.1.5 Other applications

Many other interesting applications of kernel machines have been made in order to analyse biological data.

- Prediction of protein-protein interaction: in (31) the main issue is to derive protein-protein interaction information starting directly from primary structure and associated data. Using a different database of known protein-protein interaction, a Support Vector Machine (SVM) was trained to recognize and predict interactions based on its primary structure and physiochemical properties.
- Identification of peptides with data coming from mass spectrometry. In (32) peptides were identified using mass spectrometry, by means of a program called SEQUEST. SVM then was used to reduce false positives.

In conclusion, there are many advantages in using machine learning techniques and, in particular, kernel machines in bioinformatics, as Table 1 below summarizes.

**Table 1:** Advantages of using Kernel Machines in bioinformatics.

| Characteristics of biomolecular data and problems | Kernel methods characteristics |
| --- | --- |
| Big dimensions of bio-technological data | Can treat large data sets |
| Many bio-molecular data have a non vectorial structure | Can treat non vectorial data (graphs, trees, strings) |
| Biological knowledge on the data is important and available and has to be taken into account in applying machine learning methods | Can include a-priori knowledge |
| Many different data available for the same biological data | Can integrate different types of data |

**Table 2:**
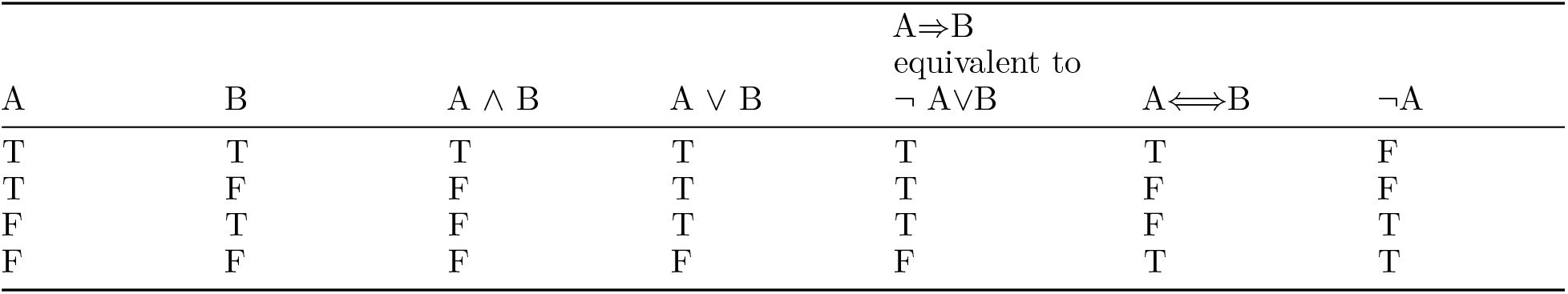
Truth table of propositional logic.

## 3 Semantic Based Regularization (SBR)

In machine learning the learning process is built upon the exchange of information between a supervisor and the learning agent. In the case of supervised learning, the learning process takes place by means of examples provided directly by the supervisor.

A large number of machine learning techniques relies on supervised or semi-supervised schemes, in which relevant interactions and therefore possible interesting information are missing. In other words there could be additional information and important prior knowledge, not taken into account in the learning process, that could be essential for optimizing the solution. Therefore to avoid finding suboptimal solutions, prior knowledge should be integrated directly into the supervised learning process.

In what follows the main idea is to represent such a prior knowledge via First Order Logic (FOL) rules, using Kernel Machines on the learning side, and posing particular attention on multi-task learning schemes. A unary predicate defined in the feature space is correlated to each task and a more abstract representation of such tasks is done by the FOL constraints. Re-stating some useful properties of kernel machines, the problem can be reformulated as the primal optimization of a function that is the composition of three terms: a loss term built on the supervised examples, a regularization term and a penalty term (coming from the constraints) (2).

This method defines a new idea of learning by kernel machines, tying logic formalism for representing human knowledge.

As described in (2) the main novelties in these methods are:

1. LEARNING FROM CONSTRAINTS IN KERNEL MACHINES: the kernel learning framework is enriched by adding the concepts of logical constraints. Unsupervised examples are used to understand to what extent the given constraints are satisfied on a random sample of data. That is why this new approach fits well with real world problems, where more importance is given to unsupervised data, due to their wide spread presence.
2. BRIDGING LOGIC AND KERNEL MACHINES: T-norms methods are used to translate from logic constraints into real valued functions, obtaining a multi-task learning problem. This way symbolic information is added to the learning problem, defining logical connections among the various tasks. Since T-norms kernel machines operate in real-numbers space, the bridging between logic and kernel machines happens in a natural way.
3. STAGE BASED LEARNING: a possible drawback in integrating logic constraints with kernel machines learning framework is that logic adds complexity to the underlying optimization problem. This is why the function to be optimized it is no longer convex and traditional optimization techniques used for SVM training are not effective. Therefore, such a complex optimization problem is solved using a stage based learning approach: in the first optimization stage only supervised examples are considered running the learning process until convergence. Only during the second phase, constraints are added to the problem. Since logic constraints and supervised examples are strictly connected, the first stage accomplishes to the minimization of the penalty term associated to the supervised data. Therefore, the constraint optimization will be more likely to start from a solution closer to the global minimum than a random initial point.

The name Semantic Based Regularization summarizes the previous concepts.

### 3.1 First order logic

Propositional logic is a formal language, based on atomic elements called propositions and on logic connectors. Connectors provide the truth value of compounded propositions, depending on the value of truth of the connected propositions. These are the available connectors:

- AND ∧
- OR ∨
- NOT ¬
- operator ⇒
- operator ⇐⇒

In particular given two propositions A and B the following truth table can be constructed, considering the above operators:

First order logic (FOL) is an extension of propositional logic to express logic for a class of objects. While propositional logic deals with simple declarative propositions, FOL additionally covers predicates and quantification. It is based on three main elements: predicates, variables and quantifiers.

- Variable: can be any object in a relevant domain. It is defined as grounded, once a specific truth value has been assigned to it.
- Predicate: is a function where the inputs are grounded variables and the output is true or false. Using the operators defined above for propositional logic, two predicates can be connected.
- Quantifiers can be of two types:
  1. **Universal quantifier** ∀ : it signifies that some proposition is true for any object.
  2. **Existential quantifier** ∃ : it means that some proposition is true for at least one object.

Consider for example three predicates *Protein*(*x*), *Enzyme*(*x*), *N on Enzyme*(*x*) stating whether a certain variable *x* is a protein, an enzyme or non-enzyme. It is possible to state a FOL clause expressing that each protein is either an enzyme or not (8):

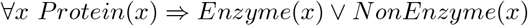

Moreover it is possible to combine variables and quantifiers. For example consider the predicate *Protein*(*x*) true if *x* is a protein and *ResidueOf* (*x, y*) true if *y* is a residue of *x*. Then the following:

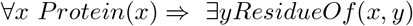

states that each protein has at least one residue.

### 3.2 An overview to Semantic-Based Regularization

Consider a multitask learning problem, with *T* being the total number of tasks and where each *k*-th task is represented by a function *f*_*k*_; therefore **f** = [*f*_1_, …, *f*_*T*_]^*′*^ defines the vector collecting all tasks function. Each function is defined in an appropriate Reproducing Kernel Hilbert Space *H*_*k*_.

In the SBR method a fundamental hypowork is that task functions are correlated since they have to satisfy a set of constraints defined by the functionals

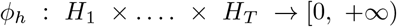

and *ϕ*_*h*_ (**f**) = 0 *h* = 1, …, *H* have to hold for any proper *f*_*k*_ ∈ *H*_*k*_, *k* = 1, …, *T*.

At this point constraints are added to the optimization function as described previously, with a new term that penalizes constraints violations, as follows:

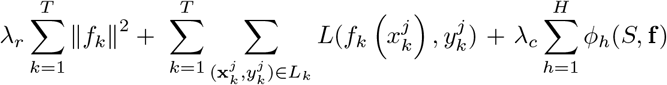

Where:

- *L* is the loss function that weights the distance of the output function from the desired one.
- *S* is the set of data points where the functions are assessed.

As shown in (2) the representer Theorem can be modified in order to show that the best solution for the above equation is stated in terms of kernel expansion.

It is possible to rewrite the previous equation using the kernel notation as:

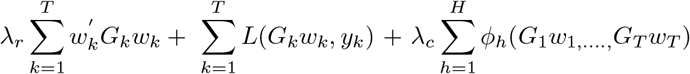

Where:

- *G*_*k*_ is the gram matrix
- *w*_*k*_ are the weights
- *f*_*k*_ are the value of the functions computed from the data sample
- *y*_*k*_ are the desired output column vectors for the patterns in a certain domain considering the *k*-th task

To optimize the parameters *w*_*k*_ of the cost function in the above equation, the gradient descent algorithm can be applied. Since *ϕ*_*h*_ in many cases are non-linear, this can represent a drawback in terms of optimization and the cost function can have many local minima. Therefore optimization can become a difficult task.

As briefly explained in the introductory section to SBR to solve this problem a two steps learning process is applied: firstly the global optimum is calculated taking all the predicates independently, posing *λ*_*c*_=0. This approach reduces the problem to a convex kernel machine one. In the second phase of the learning procedure constraints are introduced and using a gradient descent algorithm the optimal solution is found.

### 3.3 The process of translating first order logic expressions into real-valued constraints

Logic clauses will be expressed using the Prenex Normal Form (PNF form). A logic formula is said to be in PNF if it is composed by a string of quantifiers (the prefix) and by a quantifier-free part (the matrix) (33). Since every formula in classical logic is equivalent to a PNF formula, there is no loss of generality in using such a convention in writing logical constraints.

For example the following formula is in PNF, having the first part constituted by quantifiers and the second part that by a quantifier-free expression:

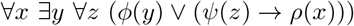

This is logically equivalent to the following that is not express in PNF:

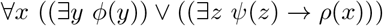

Of course the quantifier free part is equal to an assertion in propositional logic, once the variables are grounded (assigned with a specific value).

So starting from the assumption of using PNF form, the process of translating first order logic into a continuous function with values [0,1] can be done using several methods.

#### T-norms

Fuzzy logic is a logic where a value between 0 and 1 indicating the value of truth can be applied to propositions (34). In fuzzy logic t-norms are widely used in order to translate from propositional expressions into real valued functions of continuous variables.

A continuous t-norm can be defined as a function *t* : [0, 1] × [0, 1] → ℝ. Moreover the function exhibits properties of continuity, commutativity, associativity, monotonicity and 1 represents the neutral element such that *t*(*a*, 1) = *a*. A t-norm is equivalent to classical logic if the variable is near to the value 0 (false) or 1 (true).

Starting from T-norms it is possible to define T-norms fuzzy logic. This is defined through the t-norm *t*(*a*_1_, *a*_2_) that represents the logic AND, while ¬*a* is given by 1 − *a*.

The T-conorm that defines the logical operator OR, is computed as

1 − *T* ((1 − *a*_1_) ; (1 − *a*_2_)) with this definition coming from an application of the De Morgan’s law: *a*_1_ ∨ *a*_2_ = ¬ (¬*a*_1_ ∧ ¬*a*_2_).

There are a number of different t-norms logic. In particular the one used in SBR is the *product t-norm* defined as:

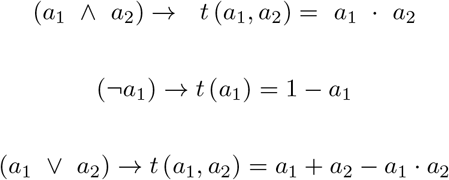

Moreover there are two possible ways to convert the operator ⇒ :

1. Modus ponens: it is possible to rewrite *a*_1_⇒*a*_2_ as ¬ *a*_1_ ∨ *a*_2_ before doing the t-norm conversion. Unfortunately this method does not encapsulate the inference process done in a fuzzy logic context (8). This is the reason why it is preferable to use the second type of conversion that is the one we used for experiments
2. Residuum: in fuzzy logic each t-norm has a corresponding binary operator ⇒ that is called residuum and that has implications. For example for a minimum t-norm the residuum is defined as:

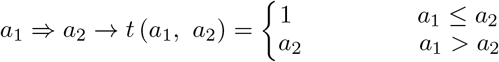

This way the condition for satisfying the implication is relaxed and the probabilistic inference problem easier and better defined.

An example regarding on how to translate implications can be the following. Consider to have the following PNF expression:

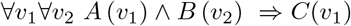

If we take the quantifier free expression and we translate it using the residuum t-norms we obtain:

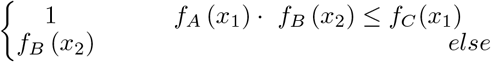

where predicates *A, B, C* have been replaced by the corresponding unknown function while the objects *x*_1_ *and x*_2_ have been represented by the grounded variables *v*_1_ and *v*_2_.

Depending on the particular type of kernel used for approximating functions *f*_*A*_ and *f*_*B*_ the representations of the objects *x*_1_ *and x*_2_ vary.

Not only operators are converted into real value ones, but also quantifiers must be converted. We will analyse separately the two quantifiers:

**Universal quantifier** ∀ : adding all the degrees of violation of the continuous expression from the t-norms considering every possible groundings for the quantified variable, we obtain the universal quantifier. For example, if the FOL formula is:

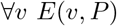

and its real valued mapping:

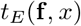

where *x* represents *v*, the conversion of the universal quantifier can be performed through the computation of the degree of non satisfaction of the expression over the domain *S* of *x*, in particular:

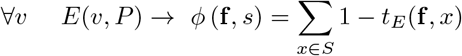

For example having the FOL formula:

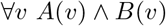

has the corresponding universal quantifier as:

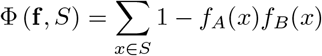

Where *f*_*A*_(*x*)*f*_*B*_(*x*) represents the t-norm generalization of *A*(*v*) ∧ *B*(*v*).

If multiple universal quantifiers are present, the conversion is recursive starting from the outer to the inner variable, that is:

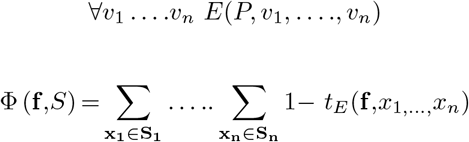

For example:

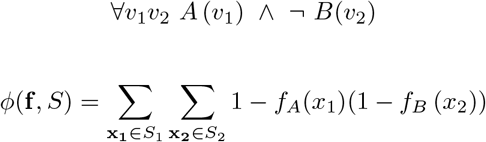

**Existential quantifier** ∃: it will be mapped in the continuous domain as

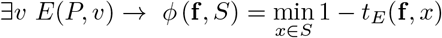

It is also possible to define the ∃_*n*_ operator, generalizing the one stated before to *n* objects:

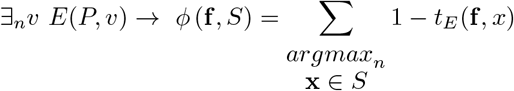

Where 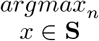 is the n assignments of *x* that maximize the value of 1 − *t*_*E*_(•) over the set **S**.

If *n* = |*S*| the ∃_*n*_ operator is equivalent to the ∀, while if *n* = 1 it is equivalent to the ∃ conversion.

## 4 Bioinformatics and biological databases

Bioinformatics plays a central role in biological research nowadays. The use of computers and machine learning tools for solving biological problems is considered extremely important for future research development. In particular information technology techniques can be applied to visualize, analyse, integrate and manage biological and genetic information. Results coming from such analysis are crucial in drug discovery research and development (35,68,69,70,71,72,73,74,75). Moreover many biological data have still to be discovered or understood and IT tools offer a wide variety of new approaches that can help in this sense. Another important role that computer science plays is that of managing and organizing biological data. A large amount of genetic and biological information has been extracted trough biological techniques and it has to be stored, waiting to be analysed. That is why computer scientists are frequently asked to search for new methods that could be useful in organizing these big databases and in building methods to implement efficient data mining techniques for extracting information.

It is possible to summarize all these bioinformatics features using the definition given by U.S. the National Institute of Health (36):

*“Research, development, or application of computational tools and approaches for expanding the use of biological, medical, behavioural or health data, including those to acquire, store, organize, archive, analyze, or visualize such data*.*”*

It is possible to analyze in details some of the main applications of bioinformatics:

- The analysis of DNA material: DNA has been at the centre of the debate since its discovery, and many different types of research occupy a central position in bioinformatics techniques. In particular DNA analysis is focused on genome sequencing and the most important genome sequencing tasks are sequence assembly, sequence similarity searching tools (for example BLAST, ClustalW), comparison between genomes, phylogenetic analysis and gene expression.
- Protein analysis: the role of proteins is crucial for every living organism and that is the reason why it is widely studied. In particular several methods have been developed for studying both the physical and the sequence structure of a protein. For the physical structure some of the most common techniques employed are X-ray crystallography, homology based models and drug design. Protein sequence is widely studied for protein family assignments, studying conserved motifs in order to understand proteins functions, proteomics data analysis and protein evolution.
- Other uses of bioinformatics: several other important uses are made of bioinformatics. In particular it is widely employed for drug designing, vaccine development, designing enzymes for detergents, genetic counselling.

As summarized by the scheme below, bioinformatics can be considered as the union of many different disciplines (35):

### 4.1 Biological databases

The vast amount of life sciences information coming as results of many scientific experiments are collected in libraries organized as biological databases. Many types of research areas produce outputs to be stored such as genomics, proteomics, metabolomics and phylogenetics (37). Therefore many different types of information can be stored in such databases such as gene function and structure, cellular and chromosomal localization, similarities among different sequences, as well as clinical effects of mutations and other genetic mutational events. Two types of databases can be identified: sequence and structure databases. The first ones are dedicated to the storage of nucleic acid and protein sequences, while the second are reserved to proteins.

The importance of these databases lies in the support that they can provide for scientists in analysing and explaining biological phenomena of different entities such as molecular interactions or the whole organism metabolism. These lines of research are fundamental, as explained before, in many different tasks such as fighting against diseases, developing of new medications and in discovering the history of life.

Each database specializes in a certain type of information. If on the one hand this type of organization is of course more efficient, on the other hand it sometimes makes it difficult to work with different databases. Biological databases are built upon the computer science concepts of relational databases, and their design and management is a core area of bioinformatics. Biological databases content is usually composed of gene sequences, textual descriptions, ontology classifications or tabular data. These type of information is referred to as semi-structured and is represented as tables, records and XML structures.

The majority of biological databases web sites offers direct access to data, and users can directly search for information online in HTML format or download it as a flat format file. Many different format types are available for data and depending on it, they are provided by different sources. For example PubMed and OMIM provide text formats, GenBank provides sequence of DNA data while UniProt sequence of protein data and, finally, protein structures are provided by PDP,SCOP and CATH.

Moreover databases can be classified into primary and derived ones. Primary databases are the ones directly built upon data extracted from experimental results, while derived or secondary databases are built using data coming from primary databases data analysis.

In particular primary databanks are the ones containing nucleotide and amino acid sequences, while the derived and specialized databanks may contain many different types of information such as genes information, protein structure, protein domains and motifs (where for protein domains are intended semi-independent regions with distinctive functions, connected to the remaining protein through a part of the polypeptide chain).

The most important primary biological databases are :

- GenBank. This database was released by the NCBI (National Centre for Biotechnology Information (39)). It contains approximately 160.000.000 of bases belonging to more than 170 million of sequences.
- EMBL datalibrary. This biological data banks was released in 1980 at the EMBL (European Molecular Biology Laboratory) in Heidelberg (Germany) (40).
- DDBJ. This is the DNA Japanese database, released by the National Institute of Genetics in Mishima in 1986. (DNA DataBase of Japan, constituted in 1986 by the National Institute of Genetics in Mishima, Japan (41).

An international agreement assures the consistency among the various DNA databases, and they are daily updated and informed about the other databases’ new releases.

Different types of organizations can provide databases. Some are released by public associations such as EMBL and NCBI while others are results of the work of academic research groups or private organizations.

Each database is constructed around an entity that is the central element around which the database is built. Each record is therefore organized containing this central element and the main attribute characterizing it.

Moreover biological databases provide a list of tools to access and manage data such as:

- Query systems tools: these are tools that can be used to perform query on databases. The most commonly used are ENTREZ, GenBank (42) query system, SRS EMBL (40) query system, DDBJ (43) query system DBGET.
- Screening tools: these tools are responsible for tasks such as sequence alignments, such as BLAST (44) and FASTA (45).
- Multiple sequences alignments tools: these tools perform alignments having multiple sequences. Examples of these tools are ClustalW, AntiClustal, ProbCons.
- Exons and regulatory elements identifications: genes are characterized by regulatory elements that allows to identify exons and genes, such as GenScan and Promoser.
- Tools for the identification of exons and regulatory elements that characterize a gene (GenScan, Promoser)

### 4.2 Protein Databanks

Many different techniques can be applied in order to obtain protein information. The protein sequence can be directly determined, translating the nucleotide sequences for which the function of the encoding gene has been predicted directly via x-ray crystallography through which it is possible to determine secondary and tertiary structure.

- UniProt/Swiss-Prot: it has been released in 1986 and it was developed in Geneva (Switzerland). Uniprot/Swiss-prot is the manually annotated and reviewed version of Uniprot (46).
- TrEMBL (*Translated EMBL)*: all the DNA sequences stored in the EMBL database have been automatic translated into amino-acid sequences and annotated as encoding proteins. This is a supplementary to SWISS-PROT.
- PIR (Protein Information Resource): this database is mainly reserved to encode protein information.
- Uniprot Consortium: TrEMBL and PIR unified, form the UniProt consortium that is a centralized repository for all protein sequences.

### 4.3 UniProt

UniProt (Universal Protein) represents the largest bioinformatics database for proteins sequences and annotation data, with many information coming from sequence genomic operations. There are three different databases constituting UniProt: [UniProt Knowledgebase (UniProtKB)](http://www.uniprot.org/help/uniprotkb), the [UniProt Reference Clusters (UniRef)](https://www.uniprot.org/help/uniref), and the [UniProt Archive (UniParc)](https://www.uniprot.org/help/uniparc).

#### UniProt Knowledgebase (UniProtKB)

(47) this database is where protein functional information is collected. Each entry in this database contains information regarding the protein such as amino acid sequence, protein name, taxonomic data, biological ontologies, classifications, as well as indication about the data source. Moreover this database is divided in two sections: a section containing manually annotated information, where information comes from curator analysis of literature and computational analysis. These two sections are referred to as UniProtKB/Swiss-Prot (the part manually annotates) and UniProtKB/TrEMBL (the part that is automatically annotated). This way, both manually and automatically annotated data is preserved, without loss of information for protein sequences. Most of the protein sequences available in UniProtKB are derived from the translating of the coding sequences (CDS) submitted to the public nucleic acid databases (EMBL-Bank/GenBank/DDBJ). All sequences coming from these databases are automatically integrated to UniProtKB/TrEMBL. All the protein sequences coming from the same gene are unified into a single UniProtKB/Swiss-Prot entry. A feature table is used to report various differences among sequence of proteins. If a protein is removed from UniprotKB/Swiss-Prot, then it is removed also from UniProtKB/TrEMBL.

#### UniProt Archive (UniParc)

(48) the majority of available protein sequences are contained in this non-redundant database. The characteristics of UniParc is to avoid redundancy, avoiding the redundancy of protein sequences that could be present in different databases and giving to each protein a unique identifier (UPI) making it possible to identify uniquely a certain protein. This UPI cannot be changed or modified. UniParc contains only protein sequences, while the rest of the information has to be retrieved from other databases using cross-references. Moreover it is possible to retrieve the history of all changes of a certain protein sequence since the various versions are stored in UniParc. Any time a sequence is changed, then an internal number is increased. Each entry in UniParc contains an identifier, the sequence, the cyclic redundancy check number, the source database with version numbers and a time stamp. Each time a new sequence is added to UniParc, it is compared with all the other sequences using the CRC64 (cyclic redundant check). If two sequences have the same value of CRC64 then these sequences will be compared as strings.

Another characteristic of UniParc is that a protein sequence is cross referenced to its origin database. In fact there is a source database identifier, a sequence identifier from the source database and a sequence version from the source database. If a sequence is deleted or changed in the source database, then the corresponding cross reference is deleted or changed. If the original database is still present, then the cross reference is useful to retrieve the original database entry, while in the case in which this is not true, then the cross reference leads to the historical sequence stored in the archive.

#### UniProt reference clusters (UniRef)

it is composed by set of sequences coming from the UniProt Knowledgebase and UniParc record, to obtain complete coverage of the sequence space at various resolutions. Sequence fragments are merged in UniRef. UniRef merges various sequence fragments. The UniRef100 database has, and mixes, identical sequences and sub-fragments with at least 11 residues from any organism into a UniRef entry. On the other hand UniRef90 clusters sequences taken from UniRef 100 with at least 11 residues and such that each cluster is composed by sequences that are identical for at least 90% sequence identity. UniRef 50 is constructed using sequences clustering UniRef90 seed sequences having at least 90 % sequence identity and 80% with the longest sequence (seed sequence) of the cluster. Further details regarding UniRef can be found in (49)

Moreover the UniProt Metagenomic and Environmental Sequences (UniMES) is a repository developed for metagenomic data. The project of UniProt was born as a collaboration between the European Bioinformatics Institute (EMBL-EBI), the SIB (Swiss Institute of Bioinformatics) and the PIR (Protein Information Resource). Many different tasks have to be performed (such as database management, software development, support) and therefore many people are involved through these different tasks. Swiss-Prot and TrEMBL were produced by EMBL-EBI and SIB, while PIR produced the Protein Sequence Database (PIR-PSD). Different protein sequences and annotation priorities exist for the two data sets. All the various part of UniProt described can be summarized in Figure 4:

**Figure 1:**
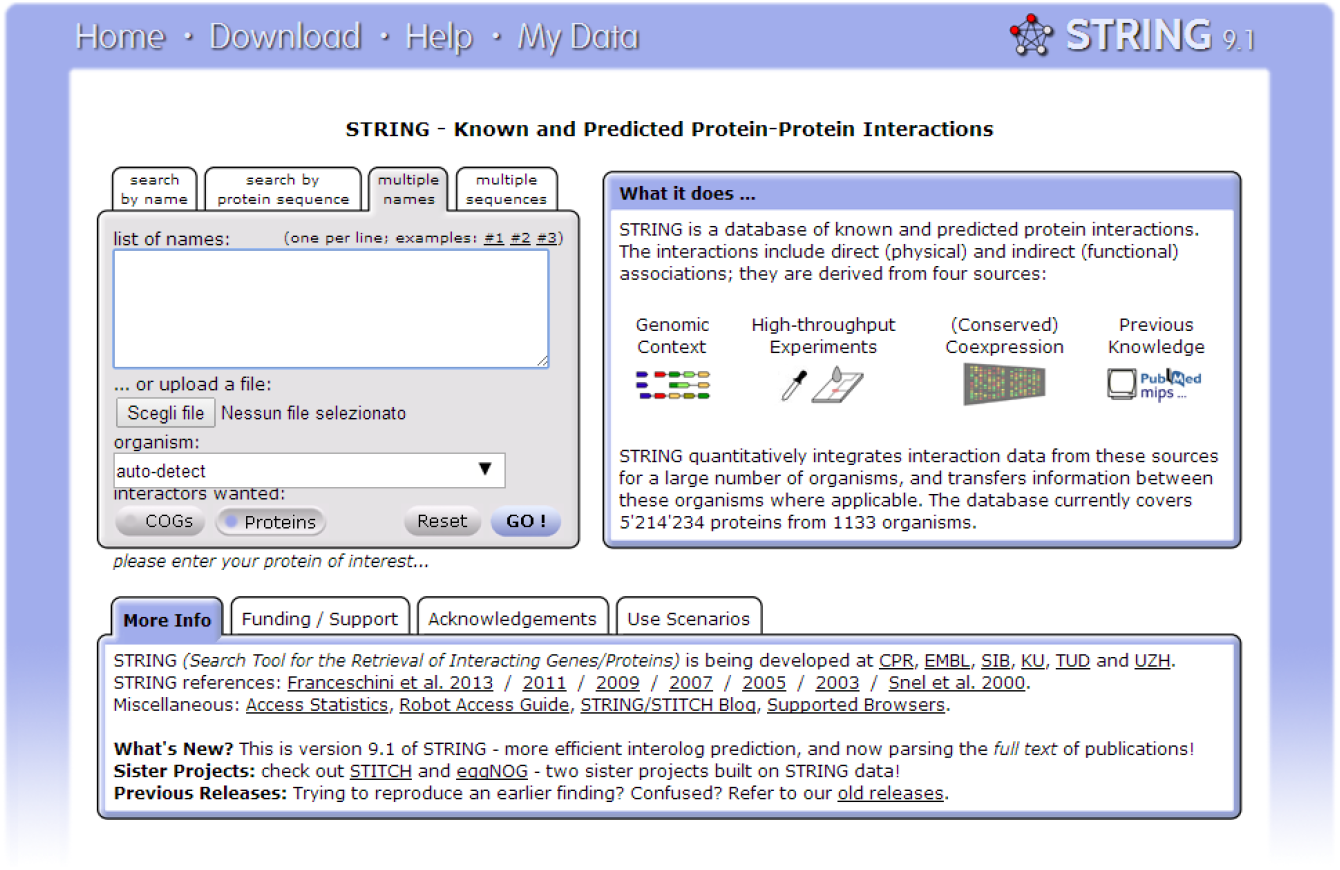
Example of the UI of STRING a protein biological database (38)

**Figure 2:**
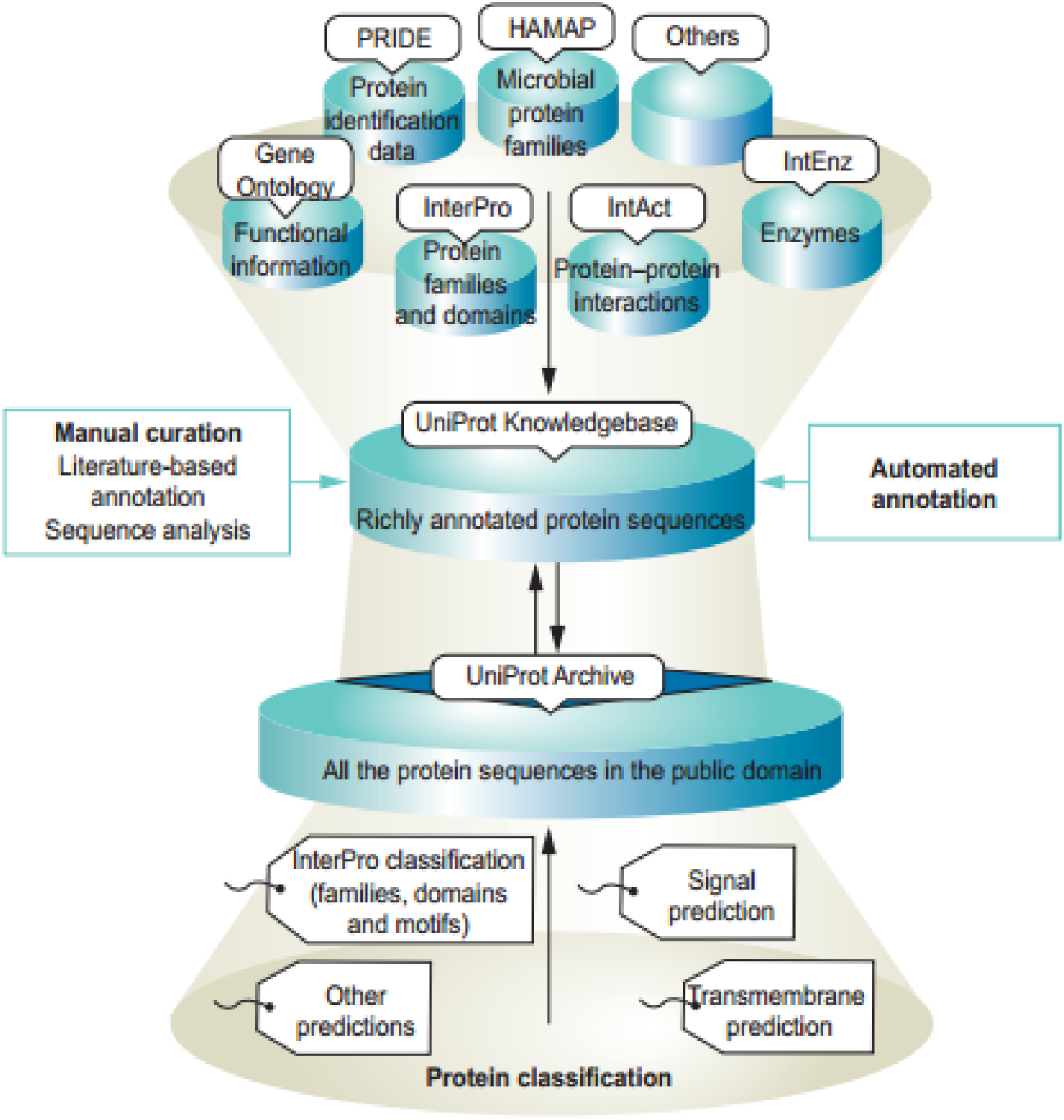
Sources of annotation for the UniProt knowledgebase

**Figure 3:**
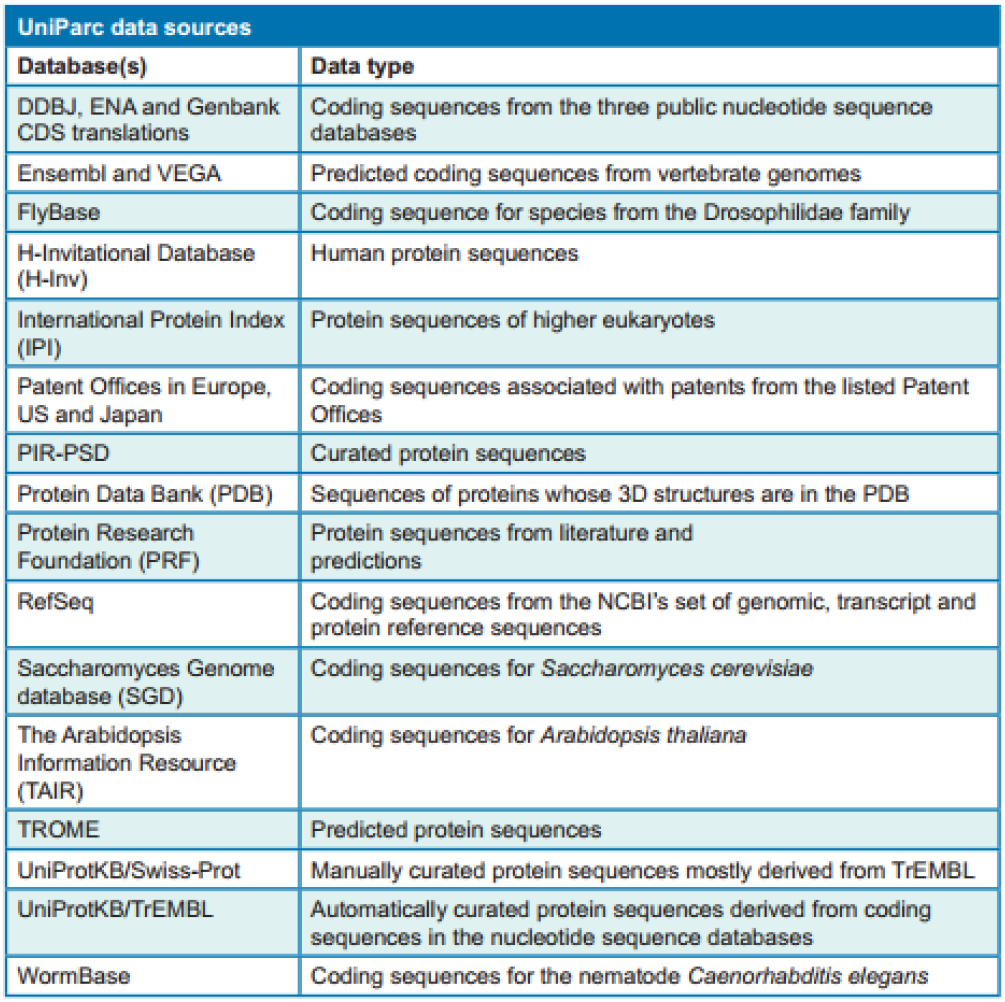
UniParc data source

**Figure 4:**
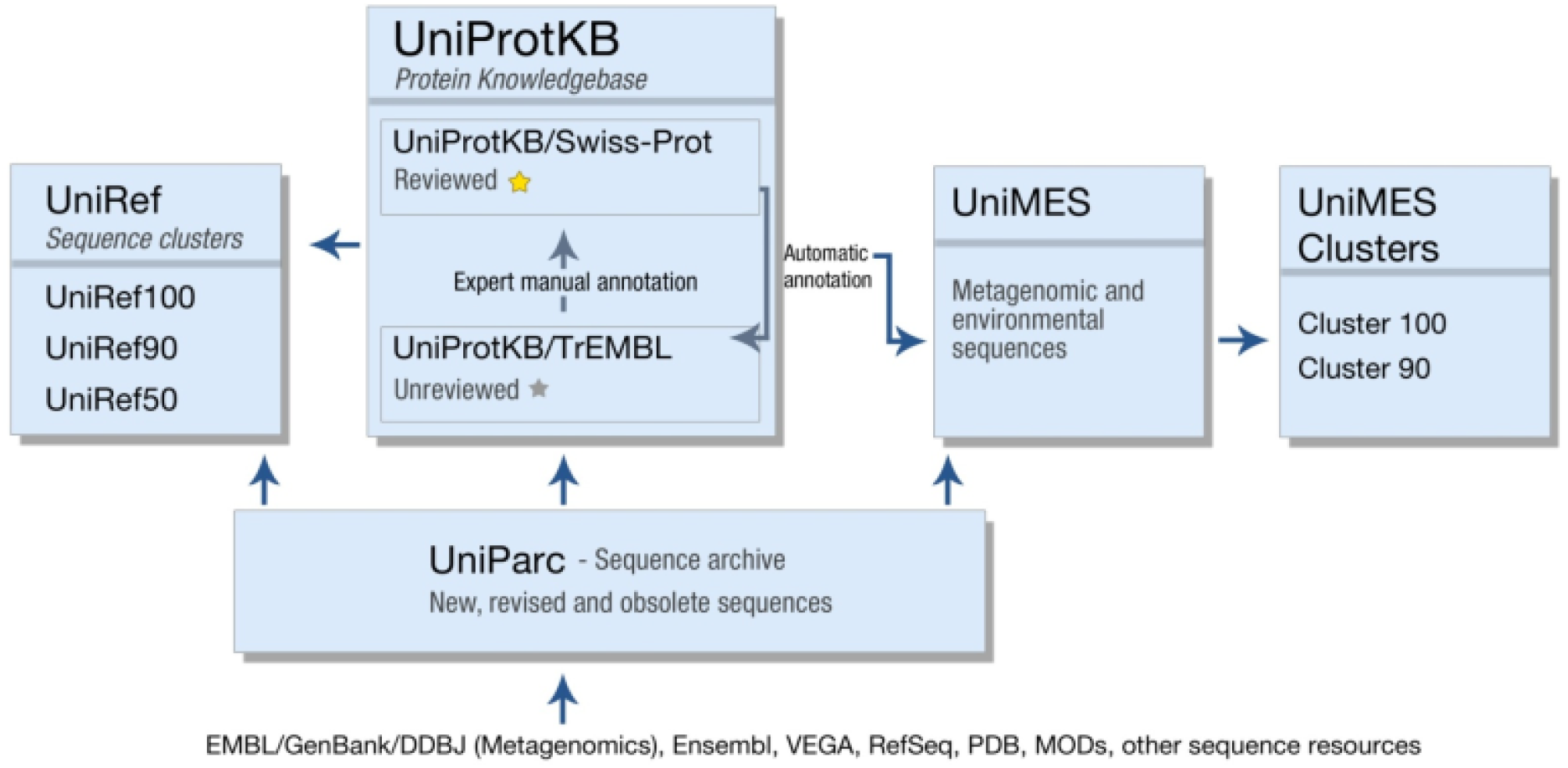
UniProt structure(50)

### 4.4 Gene Ontology

Ontologies are characterized by a hierarchical data structure that contains all relevant entities, relations among them, rules and specific constraints of the various domains. Gene Ontology (GO) was born upon the need to have constituent descriptions of gene products in various databases (51). The origin of the project goes back to 1998 and started as a collaboration among three model organism databases: FlyBase (Drosophila), Saccharomyces Genome Database (SGD) and the Mouse Genome Database (MGD). Many other databases have been included since then a large number of genomes have been included to the ontology, such as plant, animal and microbial genomes. Moreover the GO project uses three structured vocabularies (ontologies) describing gene products according to their biological processes, cellular components and molecular functions. The following graph summarizes the characteristics of each of the three domains.As an example to understand the role of the three different domains, let’s see Figure 5. This figure make an example of cellular components, molecular functions or biological processes that can be found in the Gene Ontology.

**Figure 5:**
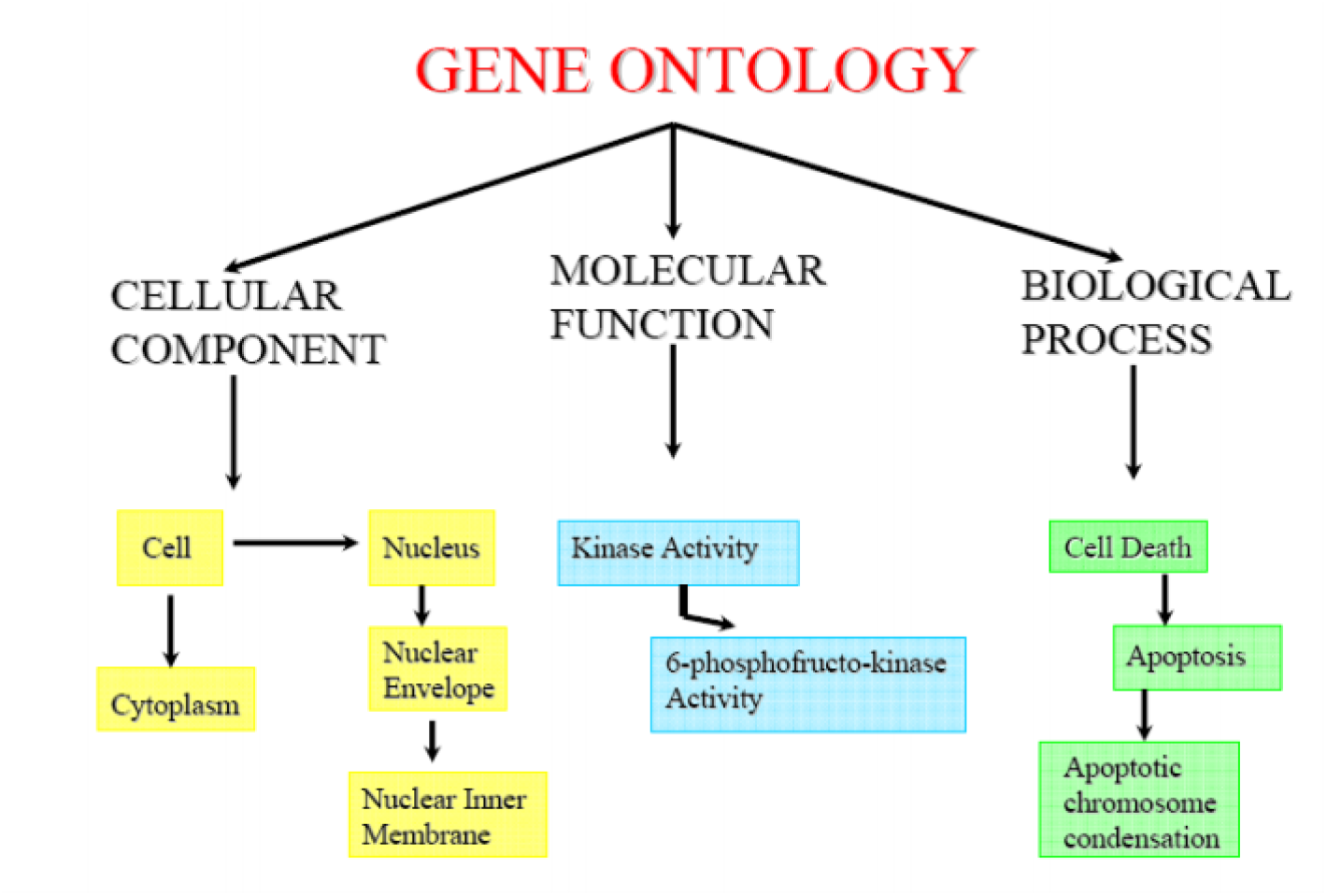
Example of the three GO domains

Three main aspects are critical to maintain Gene Ontology. Firstly the maintenance of the ontologies themselves is a difficult task, but critical for the development of the project itself. Secondly there is a continuous effort in the annotation of gene products, since it implies making association between the ontologies and the genes and gene products in the databases. Thirdly it is important to create and develop tools that maintain and create ontologies.

GO is built in a hierarchical manner, and this facilitates the task of searching for specific information. For example GO can be used to look for all the signal transduction gene products in the mouse genome, or the research can be zoomed in all the receptor tyrosine kinases. Moreover, using this, structure genes and gene products can be annotated at different levels easily.

Each GO term within the ontology has some characteristic features such as a word or a string of words, an alphanumeric id, a definition with its references and a namespace that indicates the domain to which that GO term belongs to (52). Since the GO vocabulary has been built regardless of the species, it includes terms and descriptions applicable to either eukariots and procariots, or to mono and multicellular organisms. Some terms can have synonyms that are classified as equivalent to the GO term.

GO ontology is structured as a DAG, direct acyclic oriented graph, and this means that a child (a more specialized GO term) can have multiple parents (less specialized domains) (53). Moreover there are two type of relations that a child term can establish with its parents:

1. **Is_a**: the is_a relation means that the child term is a subclass of the parent term. This type of relation is transitive; that is, consider any three GO terms *x, y, z*. If *x* is a subclass of *y* and *y* is a subclass of *z*, then *x* is also a subclass of *z*.
2. **Part_of:** this relationship means that a child GO term is always a part of a parent and in conjunction with others children it is possible to obtain the parent itself. Also this relationship is transitive. Therefore if there are three GO terms *x, y, z*: if *x* is part_of *y* and *y* is part_of *z*, then *x* is part_of *z*.

For example consider the DAG below where can be seen either is_a and part_of relationship:

The GO ontology is not static, with modifications and corrections being suggested and encouraged from the project members. Modifications are examined by the project committee and implemented if considered necessary.

## 5 Previous work on protein function prediction and dataset generation

Protein function prediction methods are widely used in bioinformatics to predict biological and biochemical functions that proteins play (54). Predicting protein functions is a fundamental task since it allows to understand proteins interaction behavior, as well as new possible biological research problems can be understood. Information on protein function may come from homologies in amino acid sequences, gene expression profiles, protein domain structures or protein-protein interaction. Many different types of functions can be played by proteins. In fact, they can be involved in biochemical reactions and transport to signal transduction. As explained in (55) protein function is a complex phenomenon that can have many different levels of action: biochemical, cellular, developmental and physiological. All these different functions can be tied together to different cellular and chemical functions and chemical functions. This is the reason why protein function should to be considered as a rather general notion, and as “everything that happens to or through a protein”.

Gene Ontology (54) is a fundamental source of information for protein functions. Querying the database with a protein name (in the Uniprot format) allows to retrieve all the information regarding cellular function, regarding the three main categories of molecular function, biological process and cellular component described in the previous section.

Biological techniques used to asses protein functions are microarray analysis, RNA interference and the yeast two-hybrid system. However these techniques are quite complex and require a long time to be performed. Since sequencing technologies have become much faster than protein function prediction techniques, this means that other methods have to be applied in order to predict protein function, to keep pace with the large amount of new sequences available. The most frequent technique for annotating sequences is done through computational methods, since they allow to perform many genes or protein annotation at once. Several different types of protein function prediction methods have been applied, and the best performing ones will be described in the following paragraphs.

### 5.1 Homology based methods

These methods are also known as homology-based function prediction. They are based on inferring functions hinging on homologous proteins whose functions are known. It is known, in fact, that proteins of similar sequences are generally homologous and therefore they have a similar function. Therefore if new proteins belonging to a newly sequenced genome have to be annotated, these new sequences can be compared with the ones of similar proteins in other genomes, in order to predict their functions. Unfortunately this type of methods are not fully reliable, since it is not always true that proteins with closely related sequences share the same functions (56). Consider for instance proteins Gal1 and Gal3 (both yeast proteins) which are an example of paralog (genes of the same genomes, that are related to a common original one, but that evolved in very different functions). In the case considered, in fact, Gal1 has become a galactokinase while Gal3 has become a transcriptional inducer (57).

### 5.2 Sequence motif-based methods

Using new protein databases where information regarding protein domains is stored, it is possible to find interesting information for protein functions identification. In fact within protein domains shorter recurrent subsequences can be found, known as motif, with which particular functions are associated. Motifs can be used in order to predict subcellular localization of a protein or other important protein functions.An example of protein families databases is Pfam (Protein Family Database) (58) a database of protein families including multiple alignments annotations. Another interesting example is PROSITE (59). This is a protein database, containing information regarding protein families, domains and other annotations including functional sites and amino acid patterns and profiles. It was developed and it is maintained by the Swiss Institute of Bioinformatics. One of its main features is the capability of discovering new protein functions analysing known protein sequences. In particular it offers tools for detecting protein sequences motifs and sequence analysis.

### 5.3 Structure based motifs

Due to the fact that 3D protein structure is generally more preserved than protein sequence, many protein functions prediction methods rely on protein 3D structure (60). Many softwares have been developed to do such comparisons among 3D protein structures.

For example FATCAT (Flexible structure AlignmenT by Chaining Aligned fragment pairs allowing Twists) is used for comparing flexible protein structures (61) automatically identifying similar 3D protein structures.

Moreover since many proteins do not have known 3D structures, there are some function prediction servers that primarily predict the 3D protein structure and then use these predicted structures to perform protein functions prediction. An example of this type of servers is RaptorX (62). An example of the 3D structure prediction is showed in the figure below:

#### Genomic context-based methods

Instead of comparing sequences for protein functions prediction, genomic context based-methods rely on the comparison between new genes/proteins and already annotated ones to make predictions regarding new protein functions (54). These types of methods are also known as phylogenomic profiles ligand and based on the observation that proteins having the same pattern (absent or present) in many different genomes usually have the same functionalities (63).

Context-based methods are important to predict cellular function or the biological process in which a certain protein plays a role. It is known in fact that proteins sharing the same signal transduction pathway usually share the same genomic context among all the species (54).

Many different types of genomic context-based methods can be analysed, that come from biological phenomena taking place inside the cell.

- Gene fusion: it happens when, through evolution, two or more genes encoding different proteins, have become one single gene in another organism.
- Co-location: chromosomal proximity can be used in order to predict functional similarity between proteins, since genes that are close in the genome usually encode proteins that have the same functions.

### 5.4 Network-based methods

Proteins or genes can be organized in a functional association network. These kind of networks are used to represent shared or similar functions within a group of proteins or genes. They are built so that each node represents genes or proteins and edges are present if there is a shared function between the two.

Consider for example figure 10:

**Figure 6:**
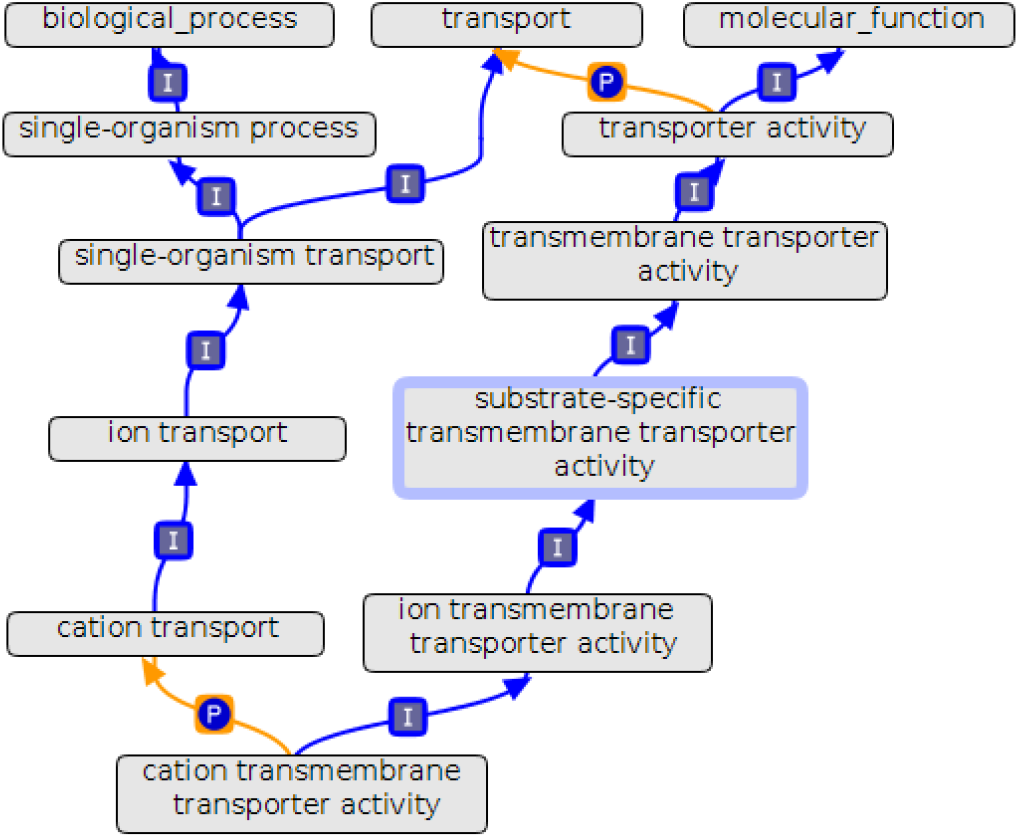
GO terms DAG

**Figure 7:**
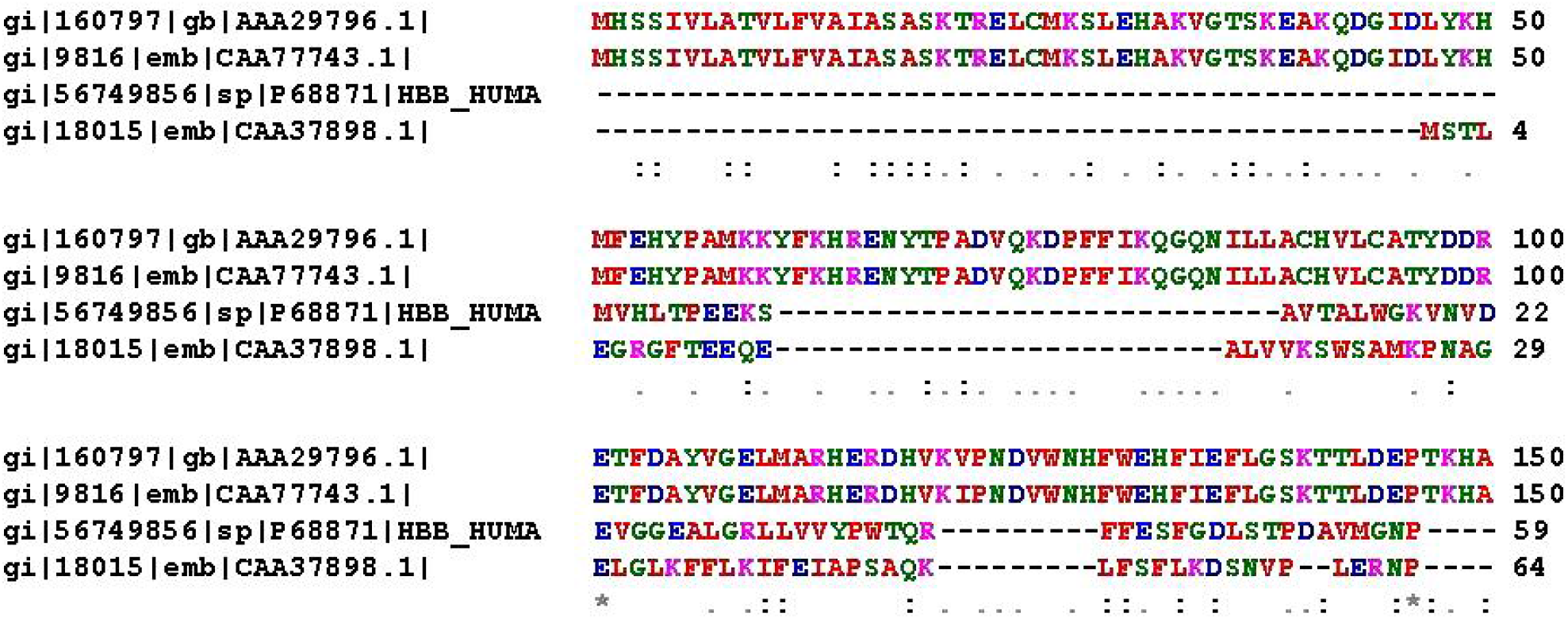
Example of a multiple alignments of 4 different haemoglobin protein sequences to detect possible sharing functions (54)

**Figure 8:**
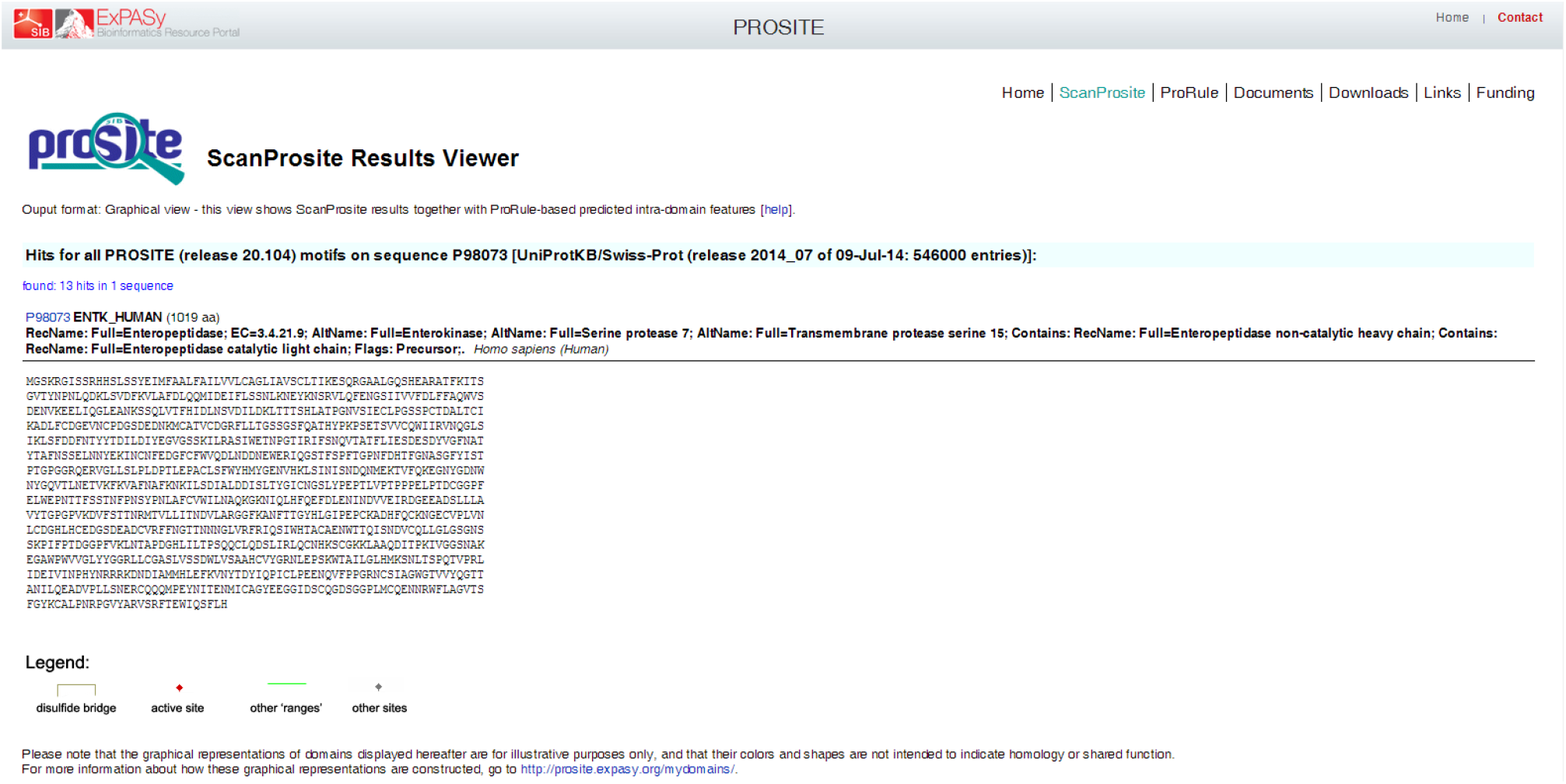
hits for all PROSITE motifs on protein sequence P98073

**Figure 9:**
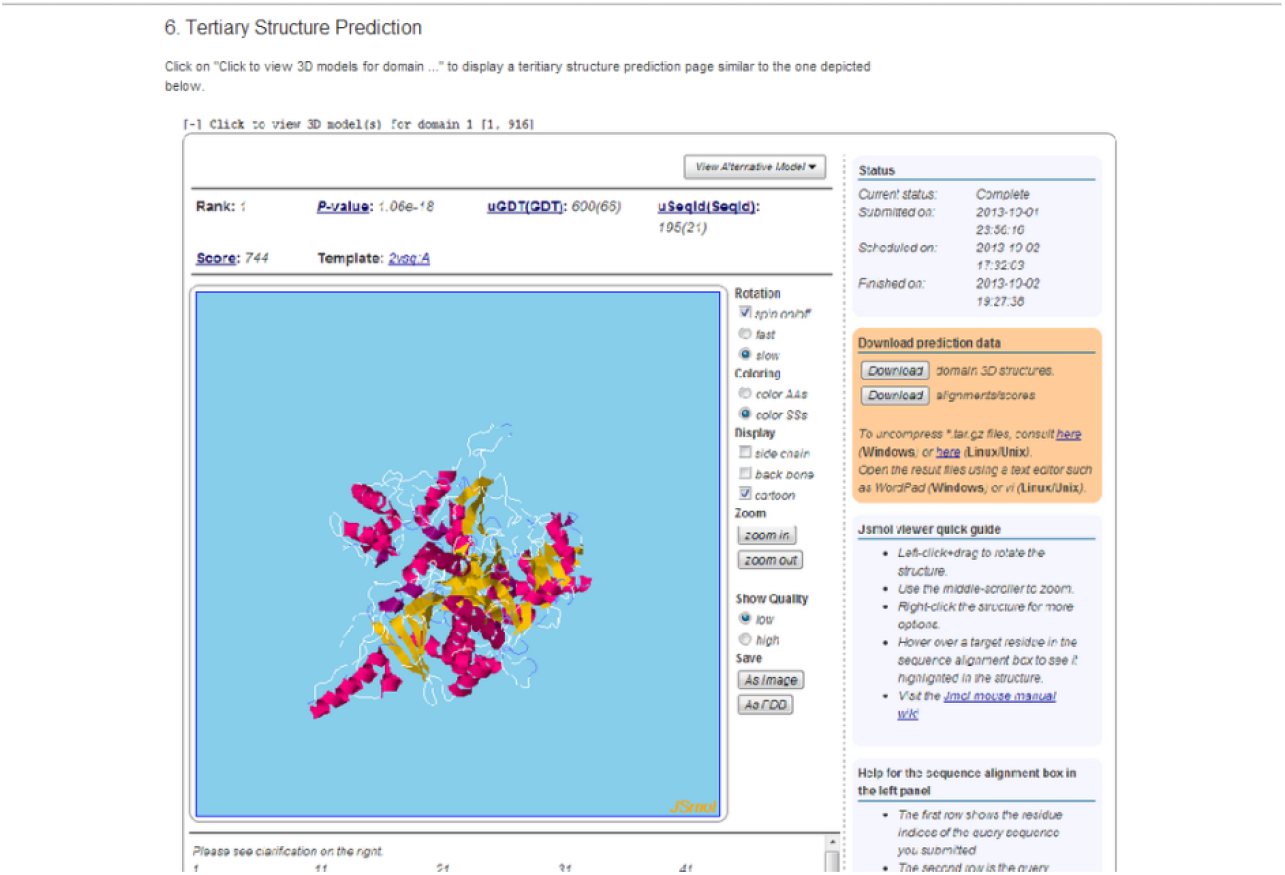
RaptorX tertiary protein prediction structure

**Figure 10:**
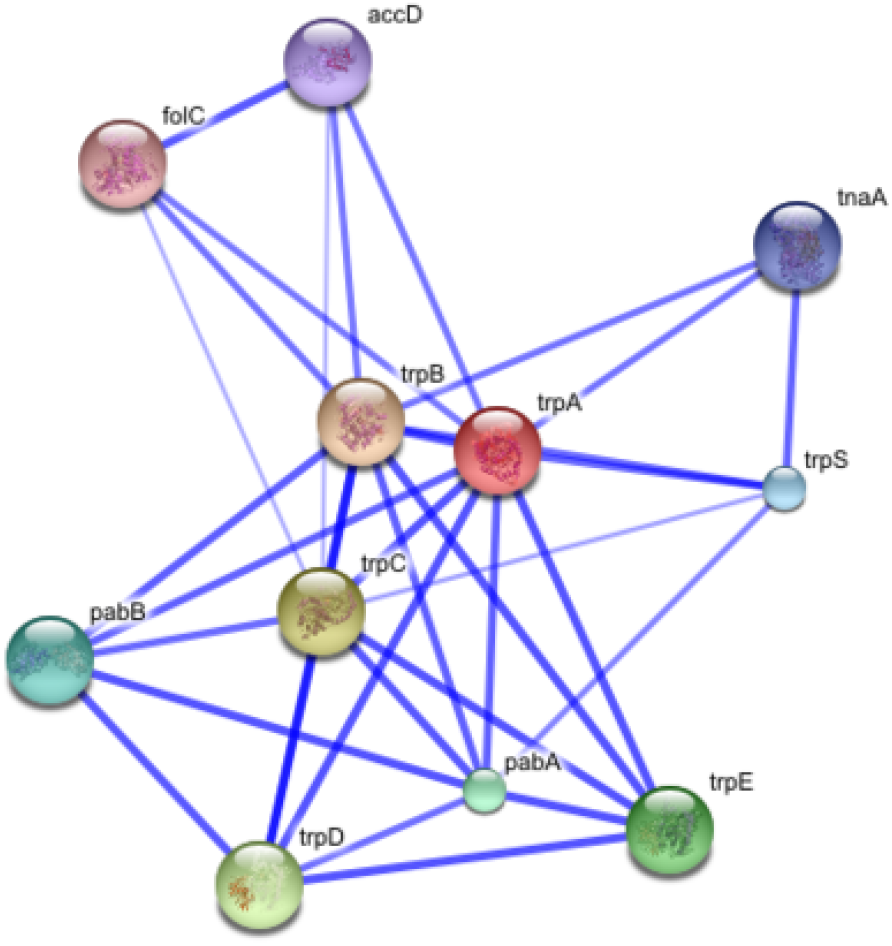
STRING interaction proteins network (54)

The figure represents a protein interaction network visualized using STRING, that is a biological database reserved to store information on protein-protein interaction (64) and that have a set of web visualization tools.

In the figure the different edges colour intensities represent the different score of confidence for a certain functional association.

### 5.5 Building the protein functions ontology for the SBR protein function prediction

In the work we performed protein functions prediction using a set of 1681 proteins that is the protein set of the yeast Saccharomices Cervisiae.

This proteins were the same used in (1). In particular in (1) the proposed method aims to apply a multi-level learning approach for the prediction of protein interaction considering three different levels: whole-protein, domains and residue, and the learning algorithm uses a multi-level learning approach as described in the following figure. In particular part (a) represented in figure 11 shows the three levels of interaction (protein, domain and residue). On the other hand part (b) shows the three different types of information flow architectures used in their article, that are the three different ways considered for exchanging information among the three levels. At last part (c) represents different ways in which two levels can be connected. For example 1 represents a way to pass training information to expand the next level training set, while 2 shows a way to pass predictions as a feature for the next level and 3 a way to pass predictions to expand training set of the next level.

**Figure 11:**
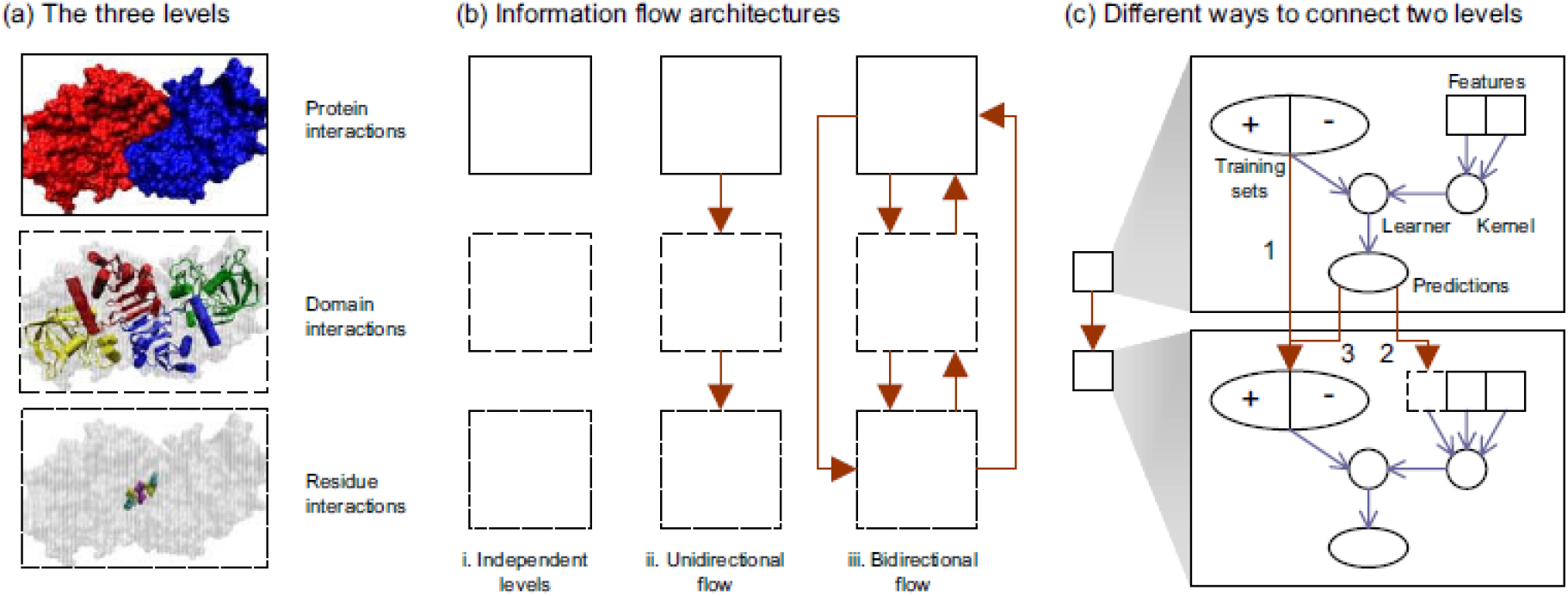
Illustration of multi-level learning concepts (1)

Also the kernel used is the protein kernel used in (1). To construct this kernel, different data features were gathered coming from a variety of sources. In particular protein features sources include phylogenetic profiles derived from COG (65), subcellular localization, cell cycle and environmental response gene expression as showed in table 3 (1).

**Table 3:** Protein Features.

| Feature | Data type | Kernel |
| --- | --- | --- |
| COG (version 7) phylogenetic profile | Binary vectors | Linear |
| Sub-cellular localization | Binary vectors | Linear |
| Cell cycle gene expression | Real vectors | Correlation (linear after standardization) |
| Environment response gene expression | Real vectors | Correlation (linear after standardization) |

To obtain the final protein gram matrix, each of these features was turned into a kernel matrix and the final one was obtained summing them.

To build the protein functions ontology for the dataset of selected 1681 proteins dataset, the following steps have been performed.

Proteins in the original dataset of (1), were named using the OLN (ordered locus name) convention, that is the nomenclature used for specifying the particular position of a gene in the genetic map. Since we needed to use information stored in the Uniprot database, the first operation was a binding operation between the OLN nomenclature and the Uniprot one. In this way the 1681 original proteins were expressed using the Swiss-Prot accession number. To perform the binding operation we referred to (66), shown in figure 12:

**Figure 12:**
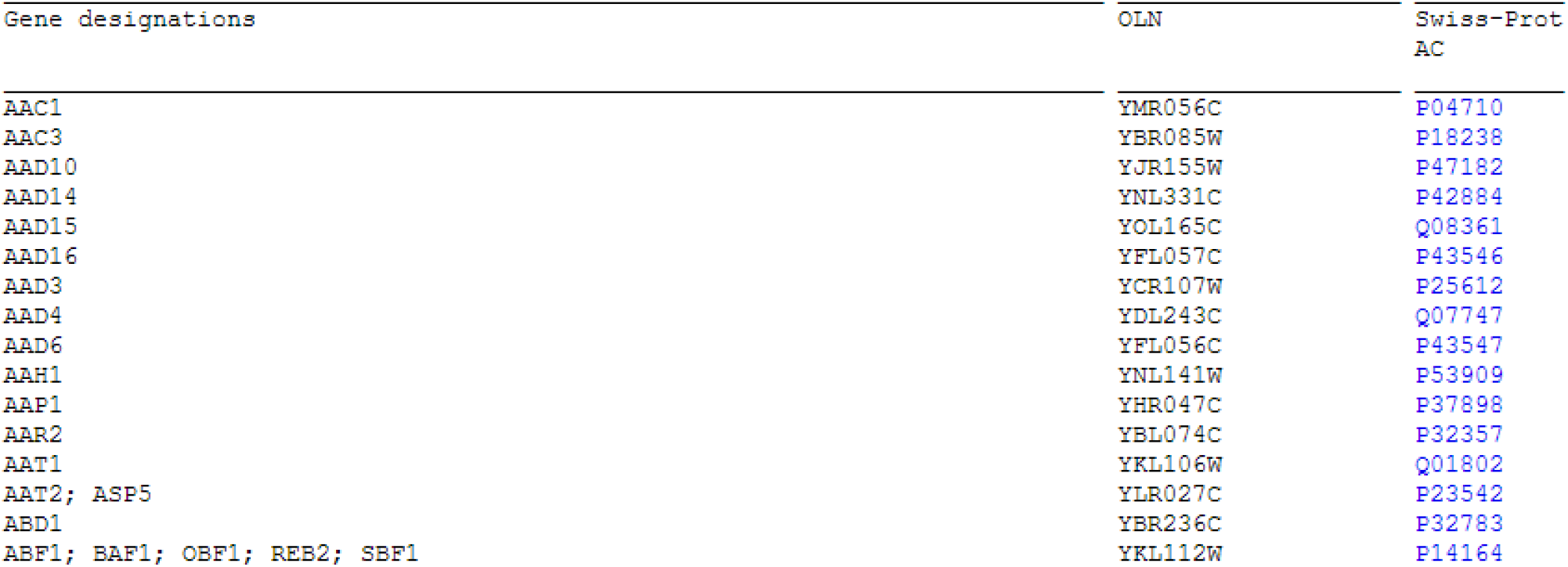
Correspondence between OLN format and Swiss-Prot nomenclature (66)

For each entry in the Uniprot database, many different types of information are stored. In particular it is available a .*txt* file where all these protein information are stored.

Among all of them there are Gene Ontology annotations, in the format shown in figure 13:

**Figure 13:**
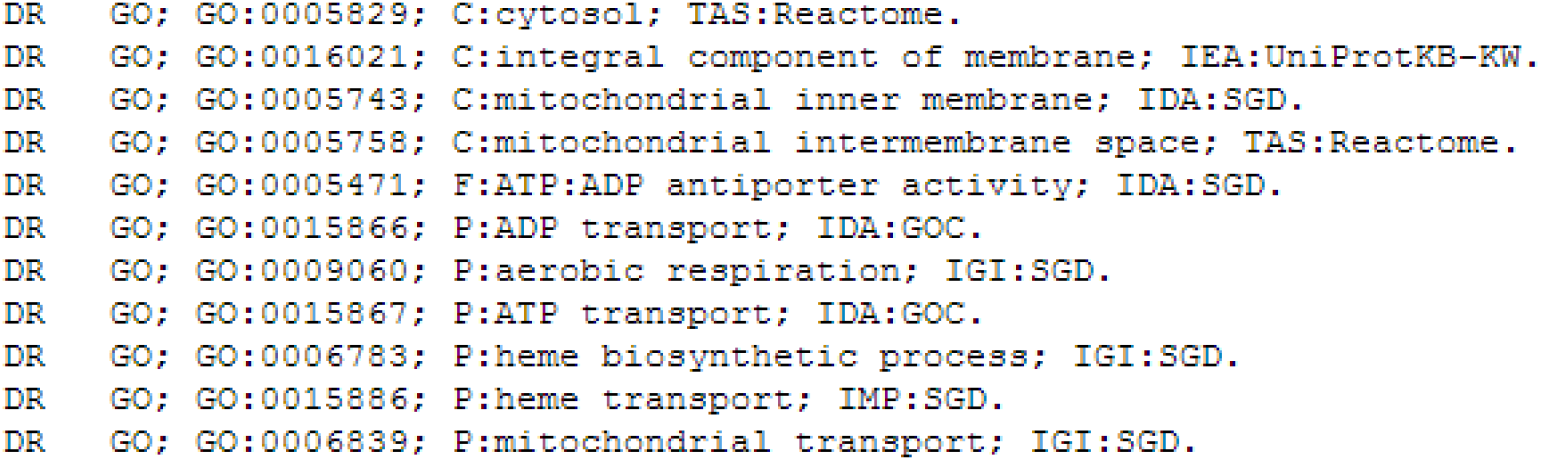
Example of the Go annotation for protein P04710

In the GO annotation each row specifies the GO entry ID node, and as we can see from figure 13 there are 3 different types of annotations (one for each ontology branch):

- C is the subcellular localization;
- F is the molecular function (that is the catalysed reaction for the enzymes);
- P is the biological process;

Since we were interested in protein interaction, we selected the biological process GO nodes for each protein, obtaining the binding between the Uniprot id term and the various GO leaves id.

In this way we obtained the whole list of GO leaves of interests. To build the protein functions ontology for each protein in the list, we started from the leaf to reach the common Biological Process root (that is 0008150) climbing up the protein functions ontology backwards. Since the GO ontology is available in various formats (67), we decided to use the OBO format. To climb up the protein functions ontology until the root we used the *is_a* relationship (previously described in the Gene Ontology section). This field is available for each term in the ontology, as shown in figure 14:

**Figure 14:**
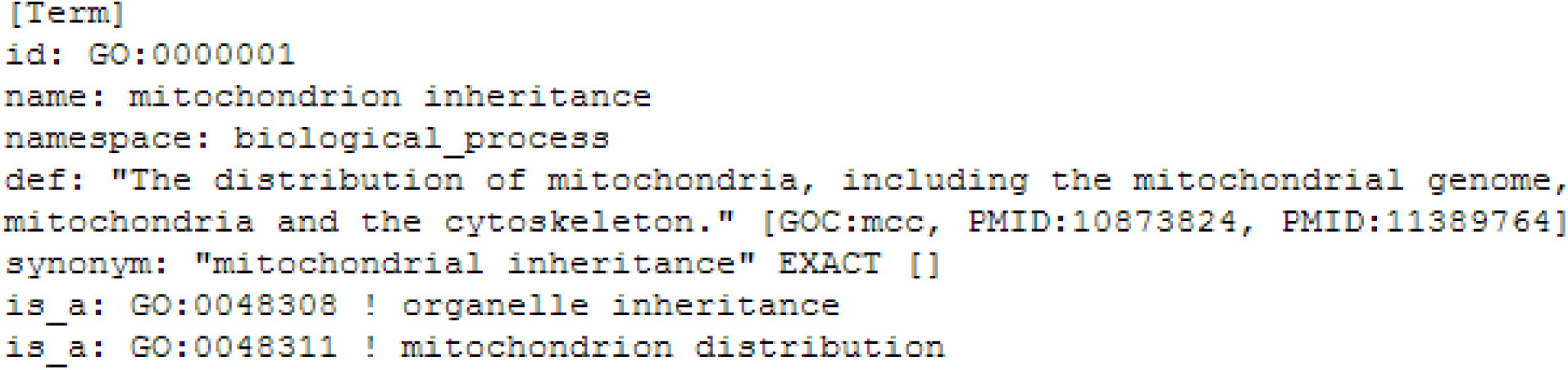
GO term 0000001

**Figure 15:**
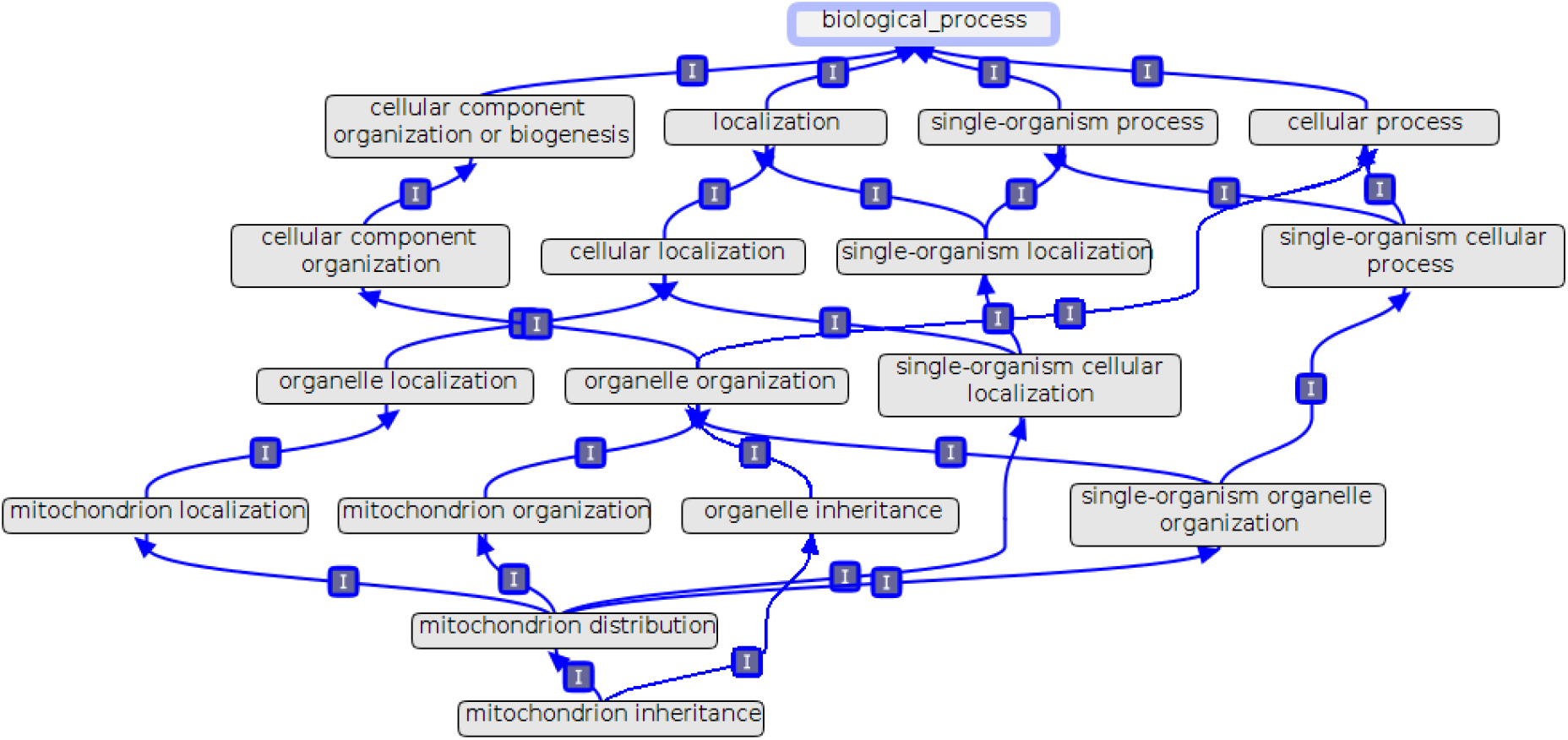
protein functions ontology

Therefore for each GO term, we obtained the corresponding protein functions ontology. An example is illustrated in the following figure:

Building this protein functions ontology was quite challenging, since as we can see from the graph there are various paths connecting the final leave with the root and they can be of different lengths.

So by considering a certain protein, and all its GO term associated with their relative trees, we were able to identify all the functions of the protein at the first/second level, considering level zero as the root level.

## 6 Experimental results

In this section we describe the experiments performed and the results obtained.

The dataset and the kernel used in this work were described in the previous section and come from (1). Since the objective for (1) was to predict protein-protein interaction using the multi-level method described in the Methods Section, to pursue our goal we had to modify the dataset.

The most important difference was in the functions to be predicted that in our case were the ones present in the protein function ontology for each protein. We concentrated on the first two levels of the trees (see figure 16) since we were interested mostly in predicting the more general classes of functions to which the proteins belonged.

**Figure 16:**
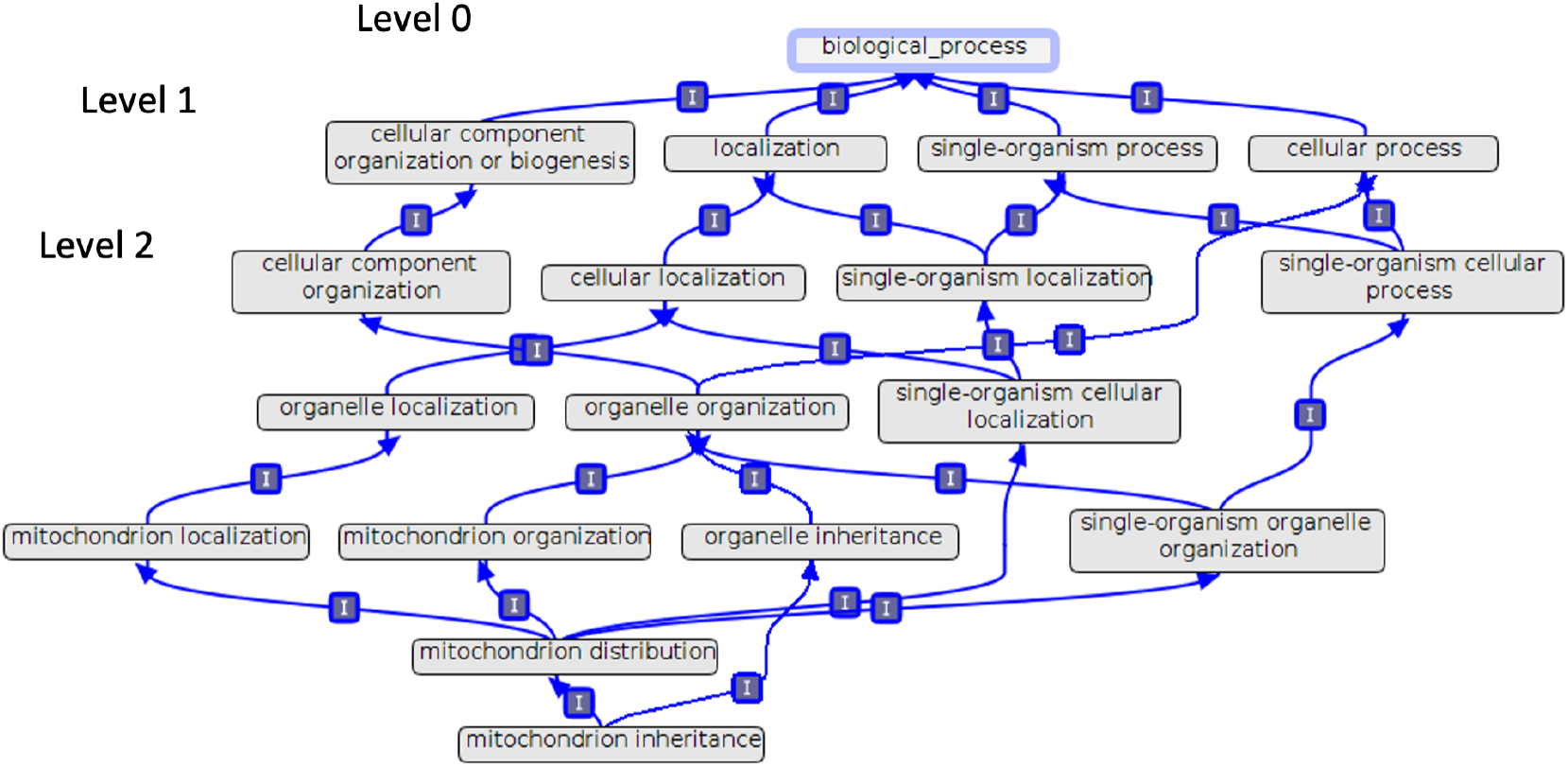
Protein Functions Ontology

Therefore to build the train and test set, and to obtain the list of all the functions to be learnt, we considered:

- each protein in the original list;
- the Gene Ontology list of nodes to which each protein is associated;
- the protein function ontology built for all the GO nodes.

In particular the functions in which a certain protein is mapped in, is the union of all the functions of level1 or of level2 of all the GO ontologies corresponding to that protein. In this way we obtained the full list of all the functions to be predicted for the first and second level, by unifying all the functions of the first and second level of each protein.

To better illustrate how the list of functions was obtained consider for example the protein YAL007C (described using the OLN name) that is part of our original list of 1681 proteins. The software we wrote to obtain the GO nodes corresponding to this protein gives as output the following:

As we can see from the figure:

- the first row shows the OLN Id of the protein considered;
- the second row corresponds to the UniProt Id, and is obtained binding the OLN id to the Uniprot one ;
- the third and the forth rows show the two id GO corresponding to that protein.

Starting from the two id GO, we built the protein functions ontologies, showed in the following figures:

As we can see from the first level in figure 17 and figure 18, the protein is mapped in the functions *establishment of localization* and *single_organism process* while in the second level the protein is mapped in *establishment of localization in cell and transport*.

**Figure 17:**
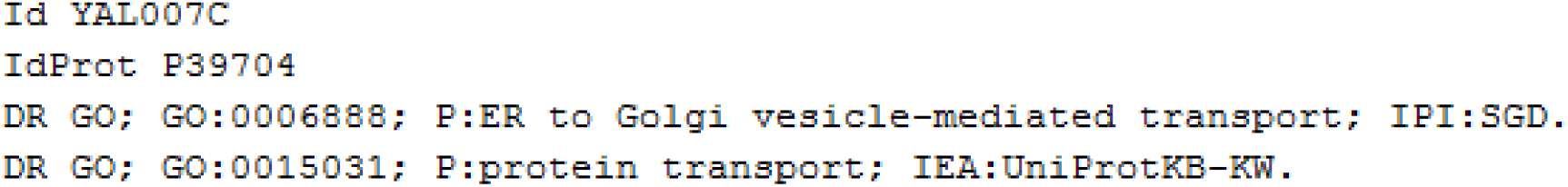
Binding from OLN to IdProt and Go ids As we can see from the figure:

**Figure 18:**
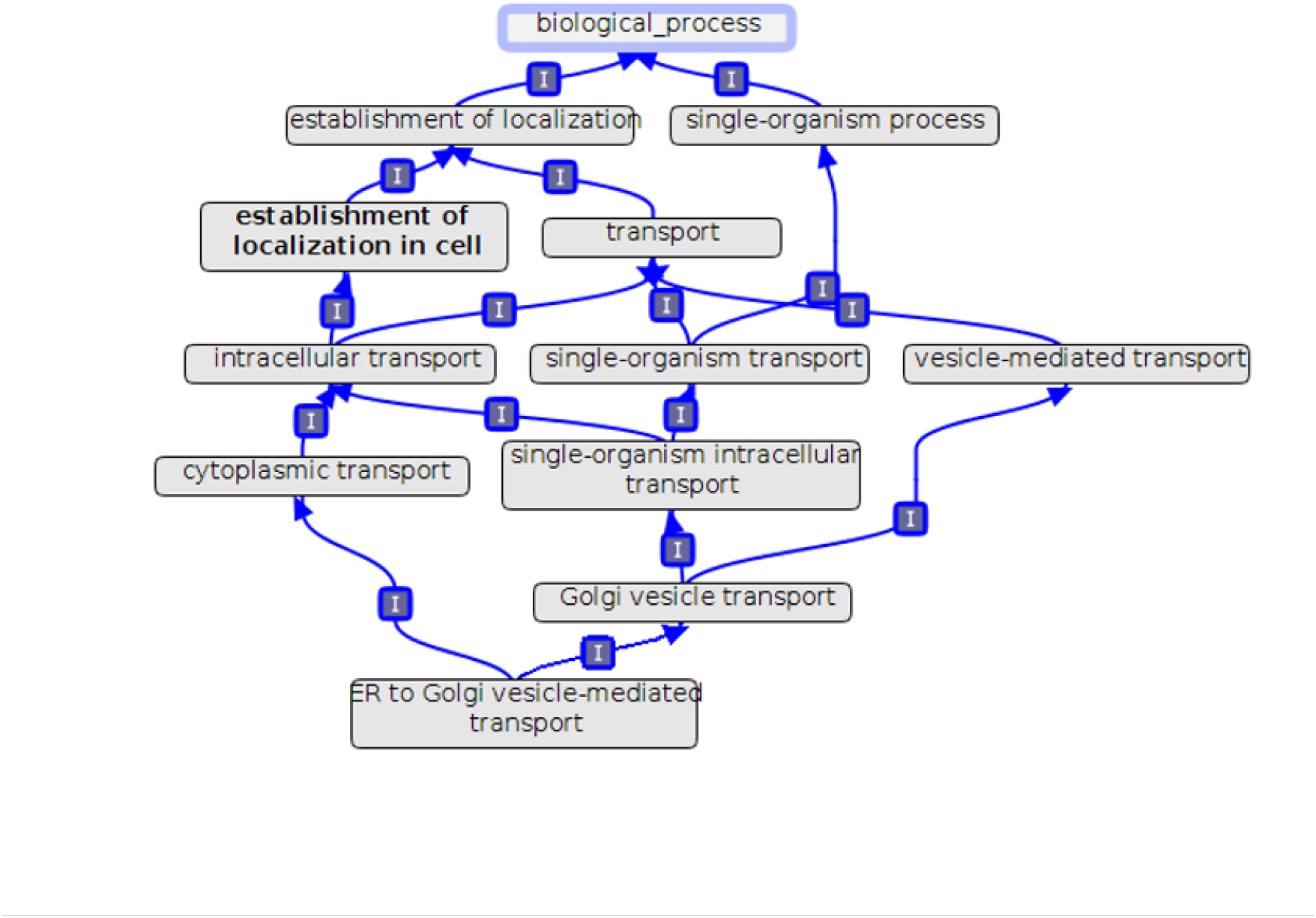
Protein functions ontology of id Go 0006888

**Figure 19:**
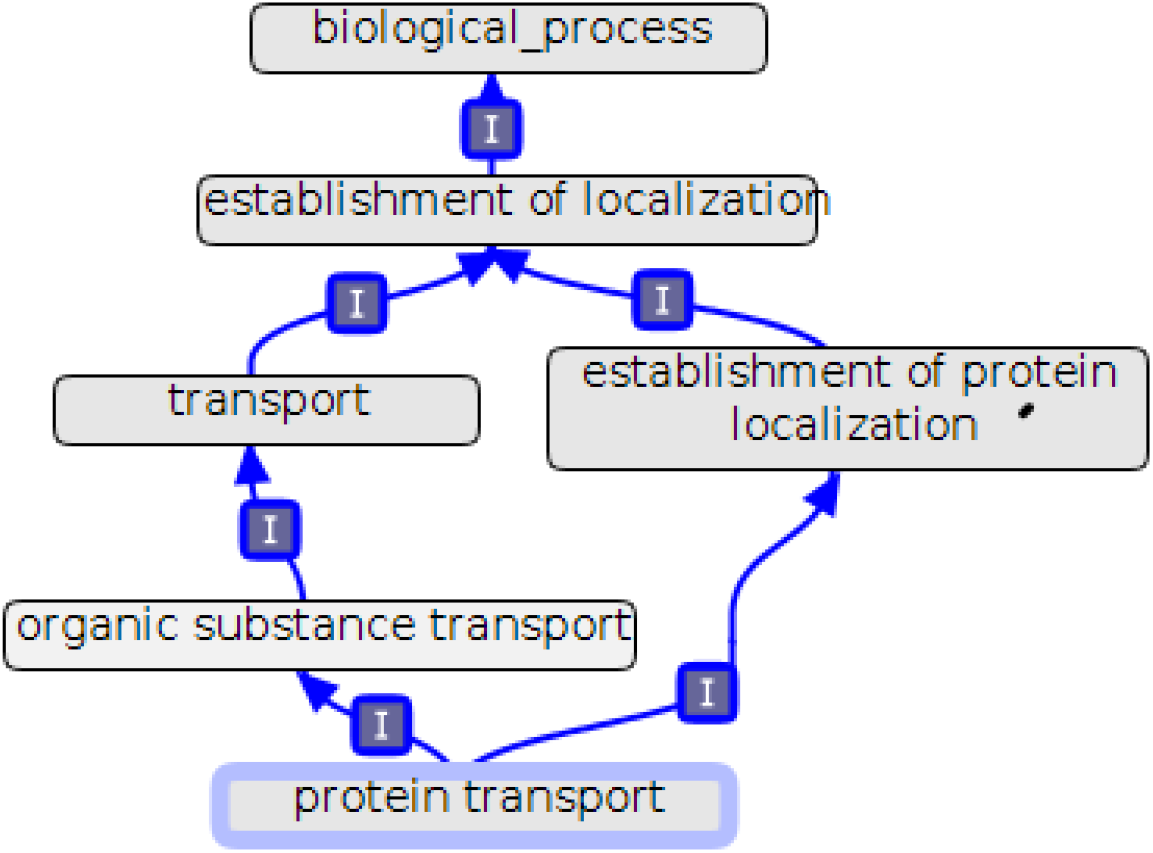
Protein functions ontology of id Go 0015031

Therefore if this was our first protein in the list, initially the list of functions of the first level to be predicted would have contained *establishment of localization and single_organism process* and the list of functions of the second level would have contained *establishment of localization in cell and transport*.

Iterating this procedure for all the proteins in the list we obtained the list of all the functions to be predicted, as well as the training and test set.

As for validation, we used a k-fold validation procedure, with *k* = 10, using the division in 10 folds already adopted by (1). Since the dataset used does not have a validation set, we chose the learning rate and the *λ*_*r*_ (the parameter controlling complexity of the model), by selecting the parameters providing the best results.

To evaluate the results we used both macro and micro statistics, in particular considering F1, recall and precision. The difference between micro and macro statistics is that while the former are obtained as an average over the single patterns, the latter ones are obtained considering the average over the results for the single classes.

For the sake of completeness, below we report the definitions of F1,recall and precision:

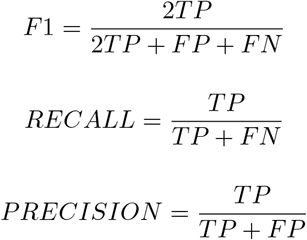

where:

- *TP* = True Positives
- *FP* = False Positives
- *TN* = True Negatives
- *FN* = False Negatives

Since as mentioned above the dataset was initially built in [1] for a different classification task, we observed that a too low number of examples was available for the functions prediction task which is of interest for this work. Therefore we decided to restrict our predictions only to the classes having at least 40 examples.

We will now describe each experiment performed, analysing and discussing results.

### 6.1 Baseline experiments

Firstly we performed experiments considering the prediction of the first and second ontology levels of functions for each protein separately and jointly.

The two levels of functions were predicted without using FOL rules. Due to this, these first set of experiments are equivalent to experiments performed in a classical kernel machine learning framework, using feature vectors only.

The rationale of these experiments was to provide a baseline for testing the prediction with SBR.

In the following table we summarize the results. The values we report are the average values obtained over the 10 folds considered:

### 6.2 SBR experiments with Ontology Consistency (OC) rules

In this set of experiments we performed joint prediction of the first and second ontology levels, using SBR as learning and collective classification tool, where we introduced FOL rules. The results were then compared with those in table 4.

**Table 4:** Kernel Machine baseline experiments.

| Ontology Levels | Macro F1 | Micro F1 | Macro Precision | Micro Precision | Macro Recall | Micro Recall |
| --- | --- | --- | --- | --- | --- | --- |
| 1 | 0.4751 | 0.4984 | 0.5093 | 0.466 | 0.588 | 0.5359 |
| 2 | 0.4521 | 0.5023 | 0.4102 | 0.4257 | 0.6497 | 0.6766 |
| 1+2 | 0.4577 | 0.4998 | 0.4378 | 0.4275 | 0.6227 | 0.6212 |

The FOL rules introduced state the dependency between functions of the first and of the second level in the Gene Ontology protein functions tree. We will refer to them from now on as OC (Ontology Consistency) rules. Overall we introduced in total 33 FOL rules.

In particular the rules used are of two types:

1. ∀*x ϵ X* : *FUNCTIONFIRST LEV EL*(*x*) ⇒ *functionsecondlevel*_1_(*x*) ∨… ∨ *functionsecondlevel*_*N*_ (*x*)
2. ∀*x* ∈ *X* : *functionsecondlevel*_*i*_(*x*) =⇒ *FUNCTIONFIRST LEV EL*(*x*)

where:

- *X* is the set of proteins
- *N* is the number of functions of the second level corresponding to a particular function of the first level

For example, consider a protein having the following protein functions ontology (for simplicity we show only the first and the second level):

Consider for example the function *localization*, that is a function belonging to the first ontology level. Then the rules stated before will be:

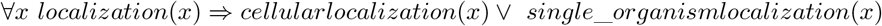

And considering the functions on the second level corresponding to localization, we will obtain two rules of the second type:

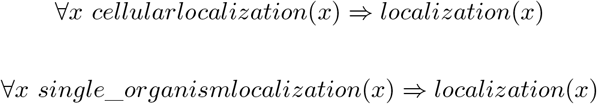

Table 5 summarizes the results obtained, comparing them with the baseline. As before the values presented are averages over the 10 folds.

**Table 5:** Table 5: Experiments comparing KM function prediction for the levels one and two of the ontology using Kernel Machines and SBR using Ontology Consistency (OC) rules.

|  | Macro F1 | Micro F1 | Macro Precision | Micro Precision | Macro Recall | Micro Recall |
| --- | --- | --- | --- | --- | --- | --- |
| Kernel Machine baseline | 0.4577 | 0.4998 | 0.4378 | 0.4275 | 0.6227 | 0.6212 |
| SBR with OC rules | 0.4875 | 0.5565 | 0.4275 | 0.4434 | 0.6773 | 0.7655 |

Results show an improvement in the performance for the case where OC rules were applied, with respect to the baseline. This suggests the ability of these rules in introducing biological knowledge that is effective for the protein functions prediction task.

Moreover taking the results of the SBR with OC rules experiments, we computed separate statistics for the predictions of first and second level of functions. In this way we could understand if the OC rules could help in improving the prediction performance of each level taken separately.

Tables 6 and 7 show the results obtained.

**Table 6:**
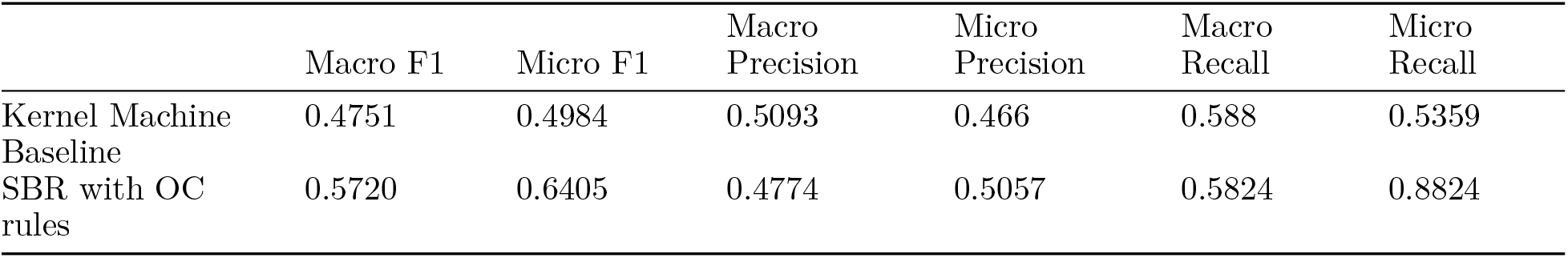
Experiments comparing KM function prediction for level one of the ontology using Kernel Machines and SBR using Ontology Consistency (OC) rules.

|  | Macro F1 | Micro F1 | Macro Precision | Micro Precision | Macro Recall | Micro Recall |
| --- | --- | --- | --- | --- | --- | --- |
| Kernel Machine Baseline | 0.4751 | 0.4984 | 0.5093 | 0.466 | 0.588 | 0.5359 |
| SBR with OC rules | 0.5720 | 0.6405 | 0.4774 | 0.5057 | 0.5824 | 0.8824 |

**Table 7:** Experiments comparing KM function prediction for level two of the ontology using Kernel Machines and SBR using Ontology Consistency (OC) rules.

|  | Macro F1 | Micro F1 | Macro Precision | Micro Precision | Macro Recall | Micro Recall |
| --- | --- | --- | --- | --- | --- | --- |
| Kernel Machine Baseline | 0.4521 | 0.5023 | 0.4102 | 0.4257 | 0.6497 | 0.6766 |
| SBR with OC rules | 0.4774 | 0.5124 | 0.4282 | 0.4300 | 0.6470 | 0.6943 |

The results confirm that the OC rules improves performances for both the first and second level of protein functions considered independently.

### 6.3 Joint prediction of protein functions and protein-protein interaction

In a real world scenario, it is often useful to classify proteins according to different aspects like: protein functions, protein localization or interactions with other proteins.

Typically, classification for a single task is performed independently. However, it is clear that independent classification does not take advantage of the fact that the different classification tasks could be correlated. For examples, two proteins can interact only if they have the same cellular localization, and proteins sharing the same functions are more likely to interact.

All this biological prior knowledge is usually neglected as no tools are available to incorporate it into the learning process. Of course this means a potential loss of information which may be useful for prediction.

The goal of these set of experiments is to demonstrate that expressing prior knowledge containing different classification tasks can be successfully exploited using Semantic Based Regularization. In particular we tied together the protein functions prediction task with the protein-protein interaction task, writing appropriate rules describing their relation.

We wrote two types of rules to describe these correlated tasks.

Let’s call the first rule PPCI1 (Protein-Protein Interaction Constraint 1). This rule expresses the prior knowledge that given a pair of proteins if they are bounded, then they tend to have the same functions.

The rule definition is:

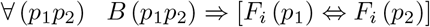

where:

- *p*_1_*p*_2_ is a pair of proteins;
- *B* is a predicate expressing whether two proteins *p*_1_and *p*_2_ interact (i.e. are bounded);
- *F*_*i*_ is any function of the first or of the second level for protein *p*_1_ or *p*_2_

The function *Bound* in this experiment is given by a supervisor from ground truth (this means that it is not learnt through the learning process). Even if this is not often realistic, it allows to establish the headroom provided by this prior knowledge.

We performed two sets of experiments with this rule, predicting jointly the first and second ontology levels.

The first type of experiments were performed using only PPCI1, while the second type of experiments were performed adding to PPCI1 the OC rules (described in the previous paragraph). This last problem is a more difficult task, considering that many more rules have to be satisfied during the learning process, adding complexity.

That is why, we tried to divide the learning process in two stages. In the first stage only the PPCI1 is used, while on the second stage OC rules are introduced. This allows the system to learn step by step, and the learning process is “simplified”. In fact, in the first step, the classification process learns only the first set of rules. Once these rules are learnt, the learning process of the second set of rules begins. All this reduces the overall complexity.

In the second set of experiments, we introduced a second type of rule expressing interaction between bounded proteins and proteins functions. We called this new rule PPIC2 (Protein-Protein Interaction Constraint 2). The definition for this new rule is the following:

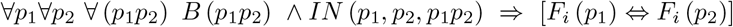

In this case the function *Bound* is not given, but has to be learnt during the learning process. This is possible by using of the function *In*, that is true if and only if the two proteins are a pair in the list of possibly interacting proteins.

Also in the case of PPIC2 the same experiments described for PPIC1 have been performed. The table 8 show the results obtained with PPIC1, PPIC2 and the baselines. Also in this case the values are the averages obtained over 10 folds.

**Table 8:** Experiments comparing KM function prediction for the levels one and two of the ontology using Kernel Machines and SBR using Ontology Consistency (OC) rules, PPIC1 (Protein-Protein Interaction Constraint 1) and PPIC2 (Protein-Protein Interaction Constraint 2)

| Rules | Number of stages | Macro F1 | Micro F1 | Macro Precision | Micro Precision | Macro Recall | Micro Recall |
| --- | --- | --- | --- | --- | --- | --- | --- |
| OC | 1 | 0.4875 | 0.5565 | 0.4275 | 0.4434 | 0.6773 | 0.7655 |
| PPIC1 | 1 | 0.5099 | 0.5718 | 0.4819 | 0.5024 | 0.6415 | 0.6448 |
| PPIC1+OC | 1 | 0.4952 | 0.5650 | 0.4282 | 0.4443 | 0.7335 | 0.7971 |
| PPIC1+OC | 2 | 0.5134 | 0.5937 | 0.4469 | 0.5137 | 0.7199 | 0.7996 |
| PPIC2 | 1 | 0.4942 | 0.5374 | 0.4716 | 0.4917 | 0.6562 | 0.6524 |
| PPIC2+OC | 1 | 0.4914 | 0.5574 | 0.4325 | 0.4490 | 0.7147 | 0.7703 |
| PPIC2+OC | 2 | 0.4977 | 0.5641 | 0.3767 | 0.4414 | 0.7411 | 0.8015 |

These results are particularly interesting.

First of all, and most importantly, they do not reject the hypowork that two interacting proteins tend to share the same functionalities, as a significant increase in accuracy is shown with respect to the learning of the first two level classes.

Moreover the two stages prediction experiments exhibit better results than the joint prediction experiments. This is confirming our working hypowork that a sequential learning process reduces complexity and therefore improves performances with respect to one-stage joint prediction experiments.

## 7 Conclusions

In this work we tackled the problem of protein functions prediction, coupled with the protein-protein interaction prediction, starting from the work of (8), using a state-of-the-art statistical relational learning technique called Semantic Based Regularization.

Semantic Based Regularization allows to express the task prior knowledge (abundant in biology) in terms of First Order Logic (FOL) clause, integrating them into the learning task.

The first step in the work was to obtain the list of protein functions to be predicted. To do so we first bounded the original list of 1681 proteins from the OLN to UniProt format. Then we created the protein functions ontology taking each GO node corresponding to each protein. Finally it was possible to create the train and the test for the various folds, considering the list of functions so obtained.

The experiments performed can be divided in two types.

In a first stage we performed protein functions prediction, while in a second stage protein functions prediction was tied together with protein-protein interaction task. In order to perform this double prediction, we wrote new FOL rules for expressing the connection between the two tasks.

The results obtained show a better performance of SBR, with respect to baseline experiments, both in predicting protein functions and in the double prediction of protein-protein interaction and protein functions.

This last task is particularly important, since showing that the two predictions can be performed jointly, can open new interesting directions to approach the PPI problem.

Moreover the joint consideration of two tasks can also help improving the prediction of a single task with respect to when they are considered separately. In particular we could incorporate information preserving proteins’ coevolution, or cellular localization, to improve protein functions prediction performances.

Further developments of this work could be to understand if there are other ways to write the FOL rules expressing such knowledge, or if alternative machine learning methods could possibly be applied to this learning task.

It would also be interesting to widen the dataset, to allow comparison with other machine learning methods.

